# Multimodal chromatin profiling using nanobody-based single-cell CUT&Tag

**DOI:** 10.1101/2022.03.08.483459

**Authors:** Marek Bartosovic, Gonçalo Castelo-Branco

**Affiliations:** Laboratory of Molecular Neurobiology, Department of Medical Biochemistry and Biophysics, Karolinska Institutet, Stockholm, Sweden; Ming Wai Lau Centre for Reparative Medicine, Stockholm node, Karolinska Institutet, 171 77 Stockholm, Sweden

## Abstract

Probing epigenomic marks such as histone modifications at a single cell level in thousands of cells has been recently enabled by technologies such as scCUT&Tag. Here we developed a multimodal and optimized iteration of scCUT&Tag called nano-CT (for nano-CUT&Tag) that allows simultaneous probing of three epigenomic modalities at single-cell resolution, using nanobody-Tn5 fusion proteins. nano-CT is compatible with starting materials as low as 25 000 cells and has significantly higher resolution than scCUT&Tag, with a 16-fold increase in the number of fragments per cells. We used nano-CT to simultaneously profile chromatin accessibility, H3K27ac and H3K27me3 in a complex tissue - juvenile mouse brain. The obtained multimodal dataset allowed for discrimination of more cell types/states that scCUT&Tag, and inference of chromatin velocity between ATAC and H3K27ac in the oligodendrocyte (OL) lineage. In addition, we used nano-CT to deconvolute H3K27me3 repressive states and infer two sequential waves of H3K27me3 repression at distinct gene modules during OL lineage progression. Thus, given its high resolution, versatility, and multimodal features, nano-CT allows unique insights in epigenetic landscapes in different biological systems at single cell level.

## Introduction

Cell identity and the underlying gene expression programs are determined through action of epigenetic modalities including transcription factor (TF) binding^1^, modifications of histones^2^, chromatin remodeling^3^, DNA methylation^4^, genome architecture^5^ and long non-coding RNAs^6^. Combinatorial effects of these factors determine the regulatory logic behind transcriptional regulation of gene expression and cell state transitions during development and in disease. It has been shown that changes in the chromatin state, for example chromatin opening^7,8^, priming by H3K4me1^9^ or enhancer RNA expression^10^ may precede gene transcription. Therefore, comprehensive chromatin maps could be used to predict cell lineage or state commitment before the transcriptional program has been activated.

The chromatin regulatory principles include co-binding of multiple epigenetic factors such as TFs, and co-occurrence of synergistic or antagonistic histone marks. For example, cis-regulatory elements (CREs) are often characterized by co-occurrence of several active histone marks^11^ and in bivalent loci, active and repressive histone marks coincide^12^. These epistatic interactions between epigenetic modalities are considered important for rapid activation and repression of the gene expression but they are still not yet fully understood.

Novel single-cell resolution technologies such as single-cell ATAC-seq (scATAC-seq) or single-cell CUT&Tag (scCUT&Tag) have provided unprecedented resolution in chromatin profiling^13–16^. These technologies enabled the detection of dynamic changes in the chromatin states in samples with unknown heterogeneity and in dynamically changing systems, such as during the development and differentiation. Furthermore, integration of single-cell epigenomics with other -omics datasets have provided additional insights into interplay between epigenetics and transcription^17,18^. However, single-cell -omics data integration relies on correlation of features across modalities, which is limiting, whereas direct measurement of multiple modalities in the same cell can bring more meaningful insights into chromatin functions.

Currently, multimodal profiling of the chromatin and the gene expression has made it possible to directly link changes in the epigenome with gene expression changes at single-cell resolution^8,19,20^. These methods provided critical insights into changes in chromatin that precede gene activation, transcription and eventually expression. Recently, technologies that profile two epigenetic modalities in single cells have been developed^21,22^. These technologies either use proteinA-Tn5 fusion pre-complexed with barcoded oligonucleotides or fusion^21^ of Tn5 to HP1α to profile active and repressive chromatin simultaneously^22^. However, these technologies either lack the flexibility to profile various modalities, have lower sensitivity, might present cross-reactivity and/or have not been used to profile complex tissues. Thus, multimodal profiling of combinations of most epigenetic modalities remains challenging.

Here, we describe nanobody-based single-cell CUT&Tag (nano-CT) that can be used to multimodally profile several epigenetic modalities in single cells. This technology allows to assay histone modifications at a single cell level with much higher depth than previous scCUT&Tag iterations^14–16,23^. We developed a novel Tn5 fusion proteins coupled with secondary nanobodies^24^ targeting either mouse or rabbit primary antibody and thus profile any combination of two histone modifications at the same time in single cells. Furthermore, we improved the CUT&Tag tagmentation and library preparation protocol so that nano-CT requires less input material and provides higher depth per cell than previous scCUT&Tag-iterations^14–16,23^. We also demonstrate that nano-CT can be combined with scATAC-seq to profile open chromatin simultaneously, altogether profiling three epigenetic modalities. The modular design of the nano-Tn5 makes it possible to profile a variety of different combinations of chromatin modalities and nano-CT will undoubtedly be an important addition to the spectrum of emerging multimodal single-cell profiling technologies.

## Results

### nano-CT improves sensitivity and has lower input requirements than scCUT&Tag

In order to multiplex the profiling of chromatin states in the same single cell, we designed a novel single chain secondary antibody^24^ - nanobody-Tn5 fusion proteins (nano-Tn5, Figure 1a), conferring the specificity of the Tn5 towards primary antibodies raised in either mouse (m-Tn5) or rabbit (r-Tn5). Because nano-Tn5 binds directly to the primary antibodies, secondary antibody incubation from the CUT&Tag protocol is not required and thus omitted. The mono-valent interaction of the nanobody with the primary antibody further allows to combine the antibody and nano-Tn5 incubation steps, greatly simplifying the nano-CT protocol and reducing the number of required washes and increasing the yield of recovered nuclei ranging from 28 to 68% (Methods, Figure 1b). This allows profiling of low input bulk samples from as little as cca. 25 000 cells as starting material, compared to previously required 200 000 cells for standard scCUT&Tag on the Chromium platform.

**Figure 1.**
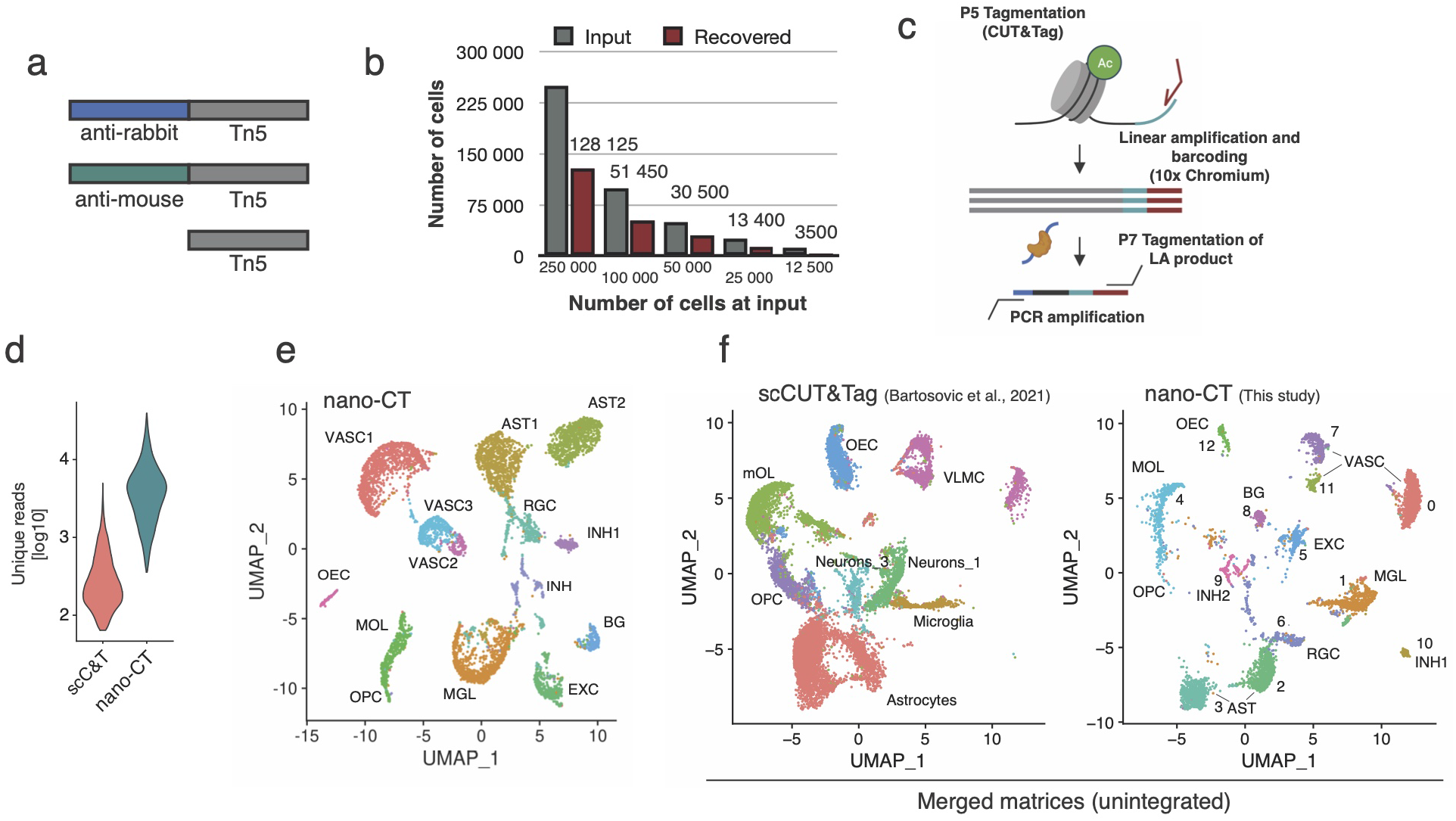
nano-CUT&Tag (nano-CT). **a** Schematic image of the Tn5 fusion proteins used in the experiments. **b**. Barplot depicting number of cells used as input for nano-CT and number of cells recovered. **c**. Cartoon depiction of the novel tagmentation and library prep strategy. The nano-Tn5 is loaded with MeA/Me-Rev oligonucleotides, tagmented gDNA is used as template for linear amplification, which is then tagmented in a second round with standard Tn5 loaded with MeB/Me-Rev oligonucleotides. The resulting library is amplified by PCR and sequenced. **d**. Violin plot depictingnumber of unique reads per cell obtained by scCUT&Tag and nano-CT using the same H3K27me3 antibody. **e**. UMAP embedding of the nano-CT data depicting the identified clusters. **f**. UMAP co-embedding of the scCUT&Tag data together with nano-CT data. Raw matrices obtained by scCUT&Tag and nano-CT were merged together and analyzed without integration. VASC – Vascular, AST – Astrocytes, RGC – radial glial cells, OEC – olfactory ensheating cells, OPC – oligodendrocyte progenitor cells, MOL – mature oligodendrocytes, MGL – microglia, EXC – excitatory neurons, INH – Inhibitory neurons, BG – bergman glia.

We further developed a novel tagmentation and nano-CT library prep protocol for single cell applications (Figure 1c). We first perform a deterministic tagmentation to capture more sites using a nano-Tn5 loaded with MeA/Me-Rev (P5) oligonucleotides. The tagmented genomic DNA is then used as a template for linear amplification and barcoding on the 10× Genomics Chromium platform. After the recovery of the barcoded DNA, a second adapter is introduced using second tagmentation with MeB/Me-Rev (P7) loaded standard Tn5, followed by library amplification by PCR (Figure 1c). This novel tagmentation protocol led to decreased cycle requirements for PCR amplification of the library in bulk, from 15 cycles to around 6 cycles (Extended data Figure 1a) yielding a library with similar concentration, although without the typical nucleosome profile (Extended data Figure 1b) The reduction in the number of PCR cycles reflects higher capture efficiency by the nano-Tn5 fusion protein and novel tagmentation protocol when compared to conventional CUT&Tag and can’t be explained solely by the introduction of linear pre-amplification step (Extended data figure 1a).

We first tested the newly developed protocol by profiling of H3K27me3 in post-natal day 19 (P19) mouse brain. We obtained 6798 single-cell profiles with a median of 3720 unique fragments, which is 15.8-fold increase in the number of unique fragments per cell over the previous generation of scCUT&Tag technology when performed on the same platform and tissue (Figure 1d). The increase in the number of unique fragments was associated with a reduction in the fraction of reads in peak regions by about 1.7-fold from 69% to 39% when using the same peak calling parameters (Extended data Figure 1d).

We then constructed a 5kb bin by cell matrix, performed clustering and dimensionality reduction using LSI and UMAP and identified 13 clusters (Figure 1e). We merged the newly generated data with the previous generation of scCUT&Tag data^15^ by combing the raw count matrices and performed dimensionality reduction (Figure 1f). The individual clusters intermingled well among the technologies (Extended data Figure 1c) without using integration methods underscoring the reproducibility of the novel nano-CT protocol with previous scCUT&Tag technology.

Importantly, we could identify and deconvolute more fine-grained clusters from the nano-CT data compared to the scCUT&Tag. For example, we deconvoluted the cluster astrocytes into three new sub-clusters (cluster 2, 3 and 6) and the cluster VLMCs into three new sub-clusters (cluster 0, 7 and 11) (Figure 1e, f). Moreover, the marker bins identified by single-cell nano-CT showed a significantly higher capture rate (23.1%) vs markers identified by scCUT&Tag (6.6%) (Extended data figure 2 a,b).

### Single-cell multimodal nano-CT in the mouse brain

Next, we loaded the nanobody-Tn5 fusion proteins with different barcoded oligonucleotides to be able to track the insertions by distinct Tn5 fusion proteins (Figure 2a). We performed nano-CT with single-cell indexing on the 10× Genomics Chromium ATAC-seq v1.1 platform on fresh mouse brain tissue obtained from 19 days old mice (P19). By using uniquely barcoded oligonucleotides, we targeted two histone modifications simultaneously, H3K27ac and H3K27me3 (2 biological replicates). In addition, we also profiled chromatin accessibility using non-fused, barcoded Tn5 (ATAC-seq) in the same sample in 2 biological replicates, altogether profiling three epigenomic modalities in single cells.

**Figure 2.**
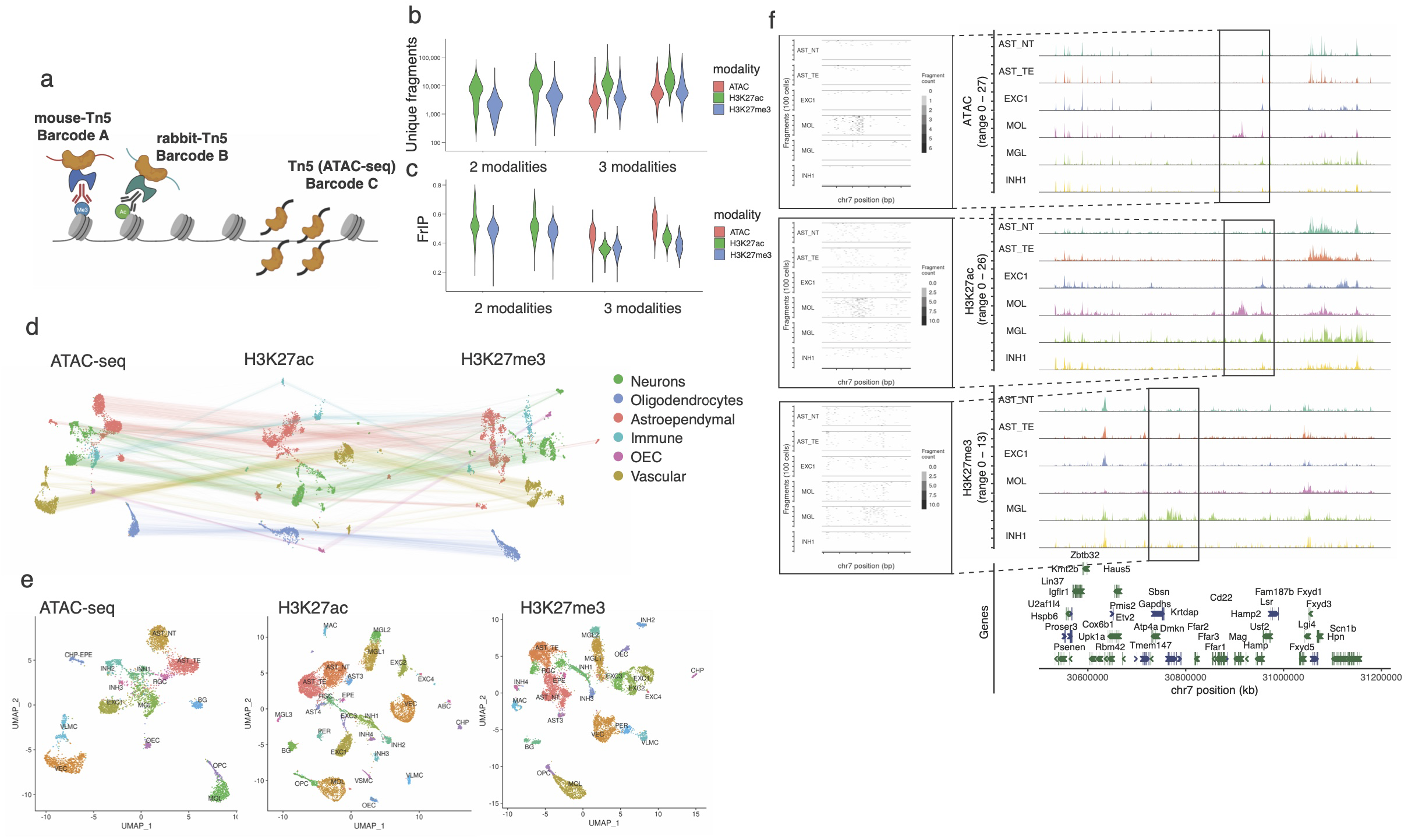
Multimodal nano-CT. **a**. Cartoon depicting the strategy used to profile multiple epigenomic modalities. Individual Tn5 and nano-Tn5 are loaded with barcoded oligonucleotides that are used in the analysis to identify the source of tagmentation and demultiplex the modalities. **b**. Violin plots depicting number of unique fragments per cell per replicate and modality. **c**. Violin plots depicting fraction of reads in peaks (FrIP) per cell per replicate and modality. **d**. UMAP embeddings of the multimodal nano-CT data for ATAC-seq, H3K27ac and H3K27me3. The lines connect representations of the same cells in the individual modalities. **e**. UMAP embedding of the individual modalities with labels of the individual clusters. **f.** Genome browser tracks of the multimodal data for several clusters showing marker peak regions. Markers – *Mag* for ATAC/H3K27ac in MOL, *Dmkn* for H3K27me3 in MGL. AST_NT – Astrocytes non-telencephalon, AST-TE – Astrocytes telencephalon, AST_3 – Astrocytes 3, AST_4 – Astrocytes 4, RGC – Radial glial cells, INH1-4 – Inhibitory neurons, EXC1-4 – Excitatory neurons, OPC – Oligodendrocyte progenitor cells, MOL – Mature oligodendrocytes, MGL1-3 – Microglia, MAC-Macrophages, VLMC – Vascular leptomeningeal cells, VEC – Vascular endothelial cells, PER – Pericytes, CHP – choroid plexus epithelial cells, EPE – ependymal cells, CHP-EPE -choroid plexus+ependymal cells, BG – Bergmann glia, VSMC – Vascular smooth muscle cells, ABC – Arachnoid bareer cells.

We demultiplexed the three modalities and preprocessed the datasets individually using the Cellranger pipeline (10× Genomics). We selected the cells using two parameters - number of reads per cell and fraction of reads in peak per cell and identified the high-quality cells using gaussian mixture model clustering in each modality separately (Extended data figure 3). Out of 13428 and 5157 cells identified in the 2-modality and 3-modality datasets 11981 (89.2%) and 4434 (85.9%) cells passed the QC filter in all modalities, respectively (Extended data figure 4a). We then compared the number of unique fragments per cell and the fraction of reads in peaks (FriP) with our previous scCUT&Tag dataset^15^. We found that multimodal nano-CT consistently outperforms previous scCUT&Tag protocol in terms of the number of fragments per cell (Figure 2b, Extended data figure 4b), with a slightly lower signal-to-noise ratio measured as the fraction of reads in peak regions (Figure 2c, Extended data figure 4c). We then constructed cell by peak matrix for all modalities and performed dimensionality reduction using latent semantic indexing (LSI), uniform manifold approximation and projection (UMAP) and clustered the cells using each modality separately. We identified individual clusters in each modality, obtaining 15 ATAC-seq clusters, 28 H3K27ac clusters and 24 H3K27me3 clusters and broadly classified them into major cell classes – Neurons, Oligodendrocytes, Astroependymal, Immune and Vascular cells (Figure 2d, 2e). We then annotated the clusters by co-embedding the active modalities (ATAC, H3K27ac) together with the single-cell RNA-seq (scRNA-seq) mouse brain atlas dataset using CCA (Extended data figure 5a, 5b). All clusters displayed a combination of unique marker regions in all modalities including enhancers labeled by H3K27ac proximal to *Mag/Mbp* (OLG), *Rbfox3* (pan-NEU), *C1qa/C1qb* (MGL/MAC), *Gad1* (INH), *Neurod1* (EXC), *Foxg1/Lhx2* (AST_TE), *Irx2* (AST_NTE) and *Foxc1* (VEC/VLMC/PER) among others (Extended data figure 6a, 6b).

In order to validate the specificity of individual de-convoluted modalities, we investigated chromatin states at the *Hox* genes clusters and found strong enrichment of H3K27me3, but not ATAC and H3K27ac, at *HoxA* locus (Figure 3a), underscoring the specificity of individual modalities at the cluster level. Similar enrichment of H3K27me3 was observed in other Hox loci including *HoxB, HoxC* and *HoxD* (Extended data figure 7a-c). Then we performed principal component analysis (PCA) of ATAC-seq, H3K27ac and H3K27me3 pseudobulk tracks for each individual cluster identified by -cell CUT&Tag^15^ (single modality) and nano-CT (multimodal). We selected the 50 most significant marker regions (peaks) from all clusters and across all modalities, merged any overlapping regions and used them as features for PCA. The PCA showed that cell populations identified through the individual modalities co-clustered together regardless of the method used for obtaining the data (Figure 3b). Then, we selected a set of highly specific peaks for both H3K27ac and H3K27me3 based on our previous scCUT&Tag data in astrocytes^15^ and generated a metagene plot of H3K27ac/H3K27me3 signal obtained by multimodal nano-CT. We observed high enrichment of the respective modifications only in the respective set of marker peaks (Figure 3c). Finally, we plotted correlation matrix of H3K27me3/H3K27ac signal in all peak regions identified in nano-CT and observed strong correlation of the respective H3K27ac and H3K27me3 tracks, and no correlation in H3K27ac-H3K27me3 combinations (Figure 3d). In summary single-cell nano-CT can be used to simultaneously obtain robust and specific multimodal epigenetic profiles of several histone modifications and open chromatin from single cells.

**Figure 3.**
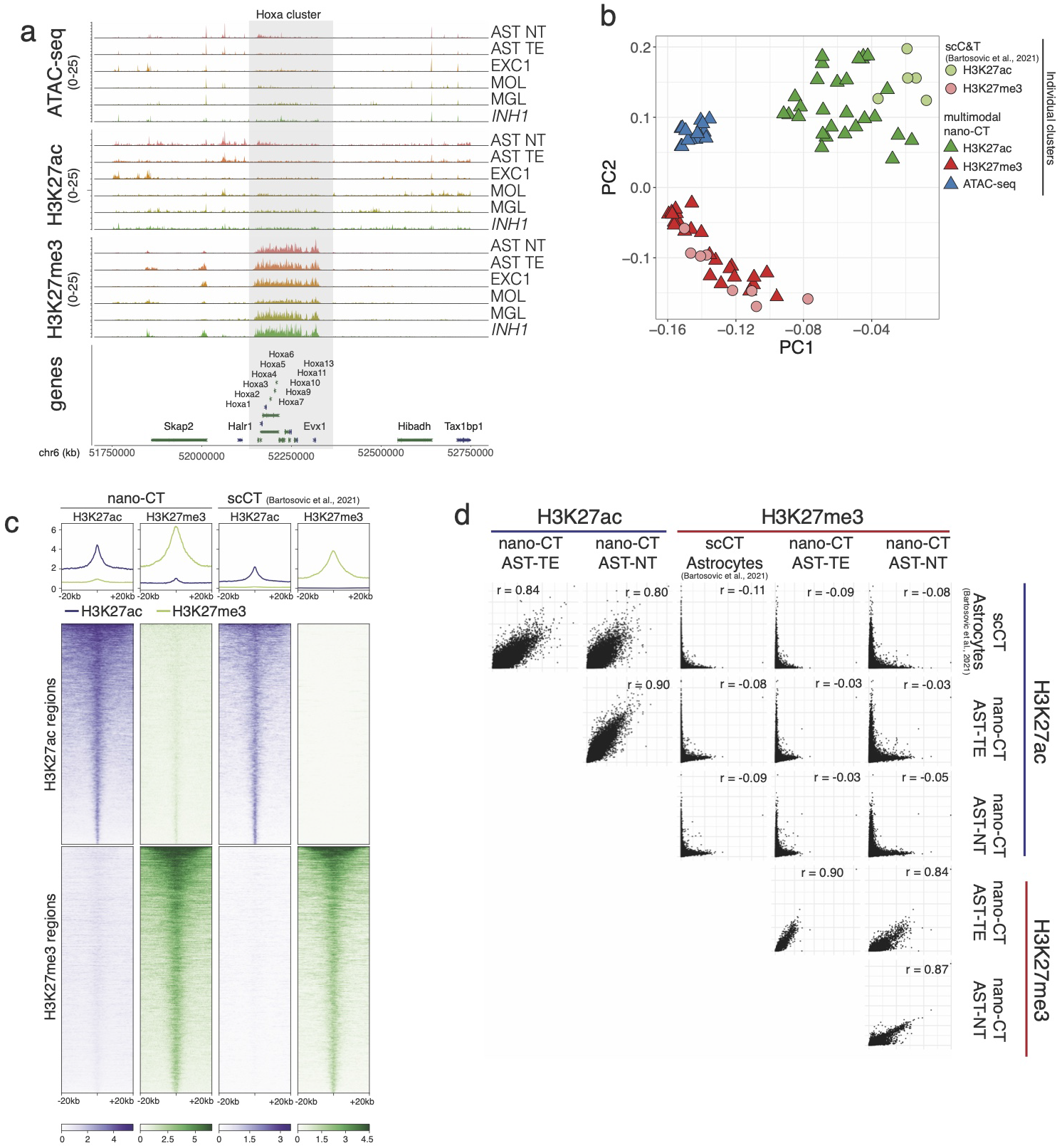
Quality control and benchmarking of nano-CT. **a.** Genome browser nano-CT pseudobulk view of the HoxA region on chromosome 6 for all three modalities. **b**. Principal component analysis (PCA) of pseudobulk tracks for each cluster identified from the respective modalities by scCUT&Tag and nano-CT. Top 50 marker regions were selected from each nano-CT cluster and modality and all peaks were merged and flattened before running PCA. **c**. Metagene plots showing the signal distribution of H3K27ac and H3K27me3 in Astrocyte populations obtained by nano-CT and scCUT&Tag around specific H3K27ac and H3K27me3 peaks. The peaks were defined and selected based on reference scCUT&Tag data^15^ **d**. Scatterplots matrix showing correlation of H3K27ac and H3K27me3 signal in astrocyte populations defined by scCUT&Tag and nano-CT. r stands for pearson correlation coefficient Cell labels as Figs. 2 and 3.

### Integrative multimodal analysis of the epigenomic state

The variability in the number of identified clusters suggests that some modalities might be more informative towards cell identity, given the similar number of fragments per cell (Figure 2b). In order to investigate whether the cell identities assigned using different modalities concord, we generated confusion matrix for all combinations of modalities (Extended data figure 8). The major cell type identities were fully recapitulated across all three modalities (Extended data figure 8a-c), whereas there were modality-specific clusters identified by the individual modalities (Extended data figure 8d-f). For example, pericytes (PER) and vascular smooth muscle cells (VSMC) can be deconvoluted from the H3K27ac modality, but not from the H3K27me3 or ATAC-seq modality (Extended data figure 8d-f).

To further improve the clustering accuracy, we used all of the multimodal matrices to perform weighted nearest neighbors (WNN) analysis^25^. The WNN largely recapitulated the clusters identified by each individual modality (Figure 4a, Extended data figure 9). We then investigated whether features that explain the most of the variability in single cells (LSI component loadings) overlapped among the different modalities. We found that H3K27ac and the ATAC-seq features overlapped the most (10601 overlapping regions), but also a large fraction of genomic regions showed variability in all three modalities simultaneously (9463) (Figure 4b). For example, the locus surrounding gene coding for transcription factor *Foxg1* was heavily regulated in two major classes of astrocytes. Whereas the chromatin surrounding Foxg1 was primarily open and K27 acetylated in telencephalon (TE) astrocytes, it was strongly K27 trimethylated in non-telencephalon (NT) astrocytes (Figure 4c). Given that Foxg1 has previously been linked with early neurogenesis in the forebrain^26^ and with inhibition of gliogenesis^27,28^ the epigenomic state of astrocytes likely reflects the developmental origin of the astrocyte populations. Other developmental and patterning transcription factors such as *Lhx2, Foxd1* or *Irx2* were also found to be differentially enriched at an epigenomic level in astrocyte populations (Figure 4d, Extended data figures 9b,c).

**Figure 4.** Multimodal analysis and visualization of the nano-CT data. **a.** UMAP embedding of the individual modalities with cluster labels identified through weighted nearest neighbors (WNN) analysis. Embedding is based on individual modalities, whereas cluster identities are assigned from WNN dimensionality reduction. **b.** Venn diagram showing the overlap of peaks identified from the individual modalities. **c, d**. UMAP projection and visualization of ATAC, H3K27ac and H3K27me3 signal intensity in single cells at the *Foxg1* **(c)** and *Irx2* loci **(d).** Grey lines connect the cells with same single-cell barcodes across the different modalities. Clusters telencephalon astrocytes (AST_TE) and non-telencephalon astrocytes (AST_NT) were selected for the visualization. Aggregated pseudobulk tracks for all modalities together with genomic annotations are shown in the right panel.

### Sequential waves of H3K27me3 deposition governs oligodendrocytes differentiation

One of the strengths of multimodal nano-CT is that it allows for direct and simultaneous analysis of dynamics of multiple histone marks and chromatin accessibility in the same cells. Our postnatal day P19 brain dataset covers the progression of the whole oligodendrocyte lineage from oligodendrocyte progenitor cells towards mature oligodendrocytes (MOLs). Therefore, we focused on the OL lineage, and we used the combined WNN embedding to generate pseudotime of oligodendrocyte differentiation (Figure 5a). We then projected the pseudotime identified from WNN onto UMAPs for the individual modalities, which recapitulated the gradient found in WNN embedding (Extended data figure 10a). Then, we investigated the magnitude of changes in all chromatin modalities at sites that are the most dynamically opening or acquiring histone modifications. We observed that loci that are opening the chromatin (ATAC-seq) acquire H3K27ac with a slight pseudotime delay, whereas there is overall little change in the overall H3K27me3 state of these sites (Figure 5b). A similar effect is observed when looking specifically at loci that acquire H3K27ac during oligodendrocytes differentiation (Extended data figure 10b). Thus, our analysis indicates that the chromatin opening precedes deposition of H3K27ac at loci that are marked for gene expression.

**Figure 5.**
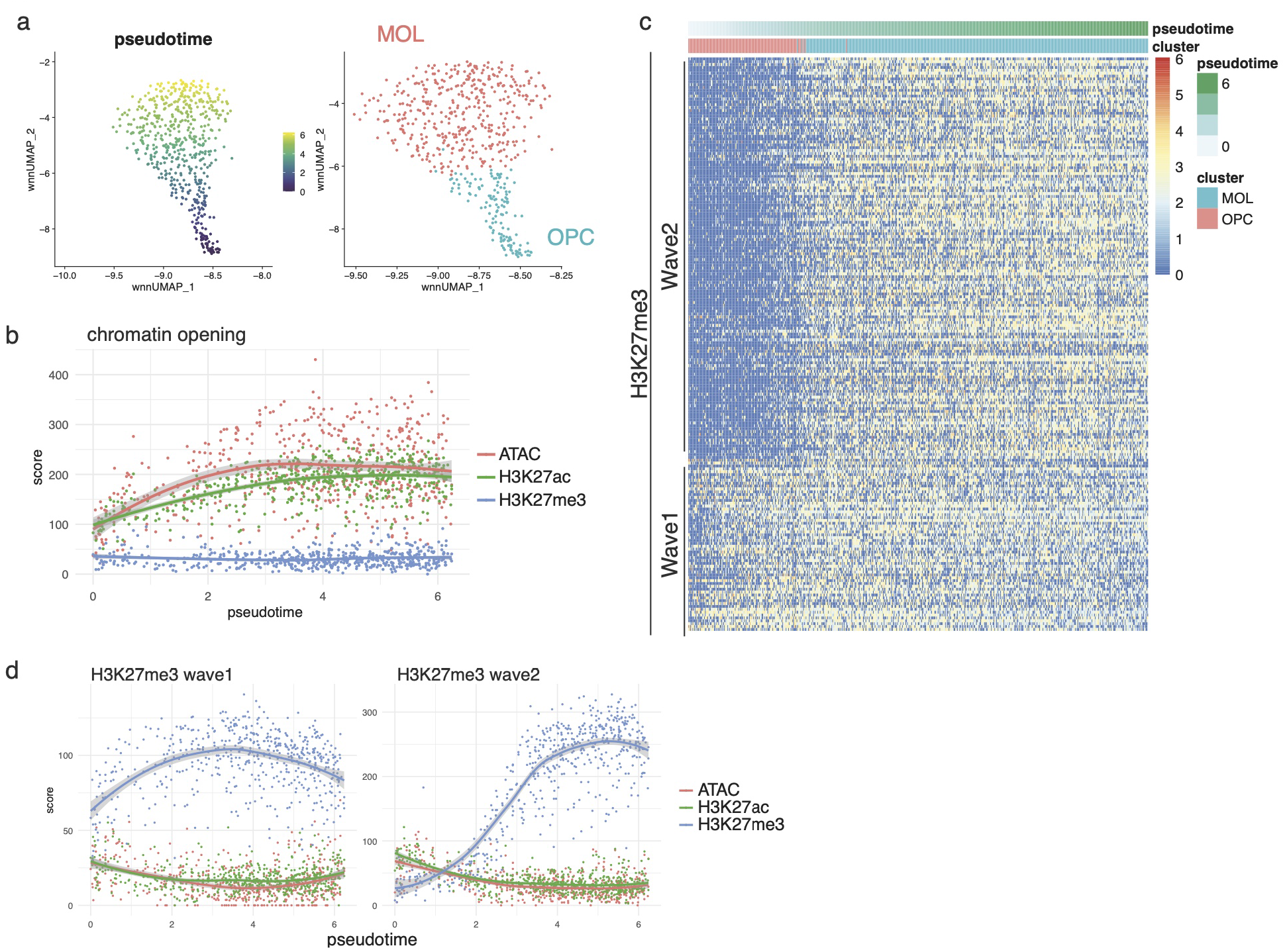
nano-CT reveals sequential H3K27me3 waves during oligodendrocyte differentiation. **a.** UMAP embedding showing pseudotime calculated by slingshot based on WNN dimensionality reduction and cluster identities. **b**. Scatter plot depicting meta-region score for all modalities (y-axis) and pseudotime (x-axis). The score was calculated as a sum of normalized score of each individual region. The regions were selected based on p-value (p<0.05, Wilcoxon test) and log fold change > 0 at the marker regions of the ATAC modality, and top 200 regions were used. **c**. Heatmap representation of the H3K27me3 signal intensity at the regions the marker regions that are gaining H3K27me3 during oligodendrocytes differentiation (p.value < 0.05, Wilcoxon test, log fold change > 0, top 200 regions). Each column depicts one single cell and row single genomic region (peak). Cells are ordered by pseudotime calculated as shown in figure 5A. The order of the regions is based on k-means clustering of the matrix with k=2. **d**. Scatter plots depicting meta-region score for all modalities (y-axis) and pseudotime (x-axis). The score was calculated as a sum of normalized score of each individual region. The regions were selected based on p-value (p<0.05, Wilcoxon test) and log fold change > 0 at the marker regions of the H3K27ac modality, and top 200 regions were used. The regions were further stratified to wave 1 and wave 2 regions based on k-means clustering as shown in figure 5c.

Strikingly, we found that H3K27me3 deposition occurs in two distinct waves during oligodendrocytes differentiation (Figure 5c,d). The first wave of H3K27me3 occurred early on in the differentiation process and was associated with repression of genes expressed predominantly in neurons, whereas the second wave repressed both neuronal genes (Extended data figure 10c) and genes expressed in oligodendrocyte progenitor cells (e.g. *Sox5, Sox6, Ptprz1*) that are associated with GO terms gliogenesis, glial cell differentiation and oligodendrocyte differentiation (Extended data figure 10d). Thus, oligodendrocyte lineage progression encompasses two sequential H3K27me3 repressive states, which would not be possible to discriminate by relying solely on transcriptiomic data. In summary, multimodal nano-CT analysis allows unique insights into the epigenomic processes driving biological processes such as oligodendrocyte differentiation.

### Chromatin velocity predicts the differentiation trajectory of oligodendrocytes

It has been shown that future transcriptional states can be predicted from scRNA-seq data using RNA velocity modeling based on the ratio of spliced/unspliced mRNA^29^. Recently, a similar concept has been proposed for chromatin velocity, using information from two anti-correlated chromatin modalities – heterochromatin and open chromatin^22^. Our multimodal chromatin profiling allows us to define several chromatin velocities – ATAC/H3K27ac, ATAC/H3K27me3 and H3K27ac/H3K27me3 velocity. Since Nano-CT indicates that chromatin opening (ATAC) precedes H3K27ac (Figure 5b), which is analogous to spliced/unspliced mRNA relationship, we hypothesized that we can directly leverage the velocity framework to predict the directionality of a differentiation pathway. We tested this idea by generating a gene by cell matrix for ATAC and H3K27ac and using these as an input into the scvelo algorithm^30^. Indeed, ATAC/H3K27ac velocity accurately predicted the differentiation trajectory of the oligodendrocyte lineage from OPCs towards mature oligodendrocytes (Figure 6a). Phase plots of ATAC-seq versus H3K27ac of several genes associated with oligodendrocytes differentiation such as *Mal* or *Mag* further confirmed this directionality (Figure 6b).

**Figure 6.**
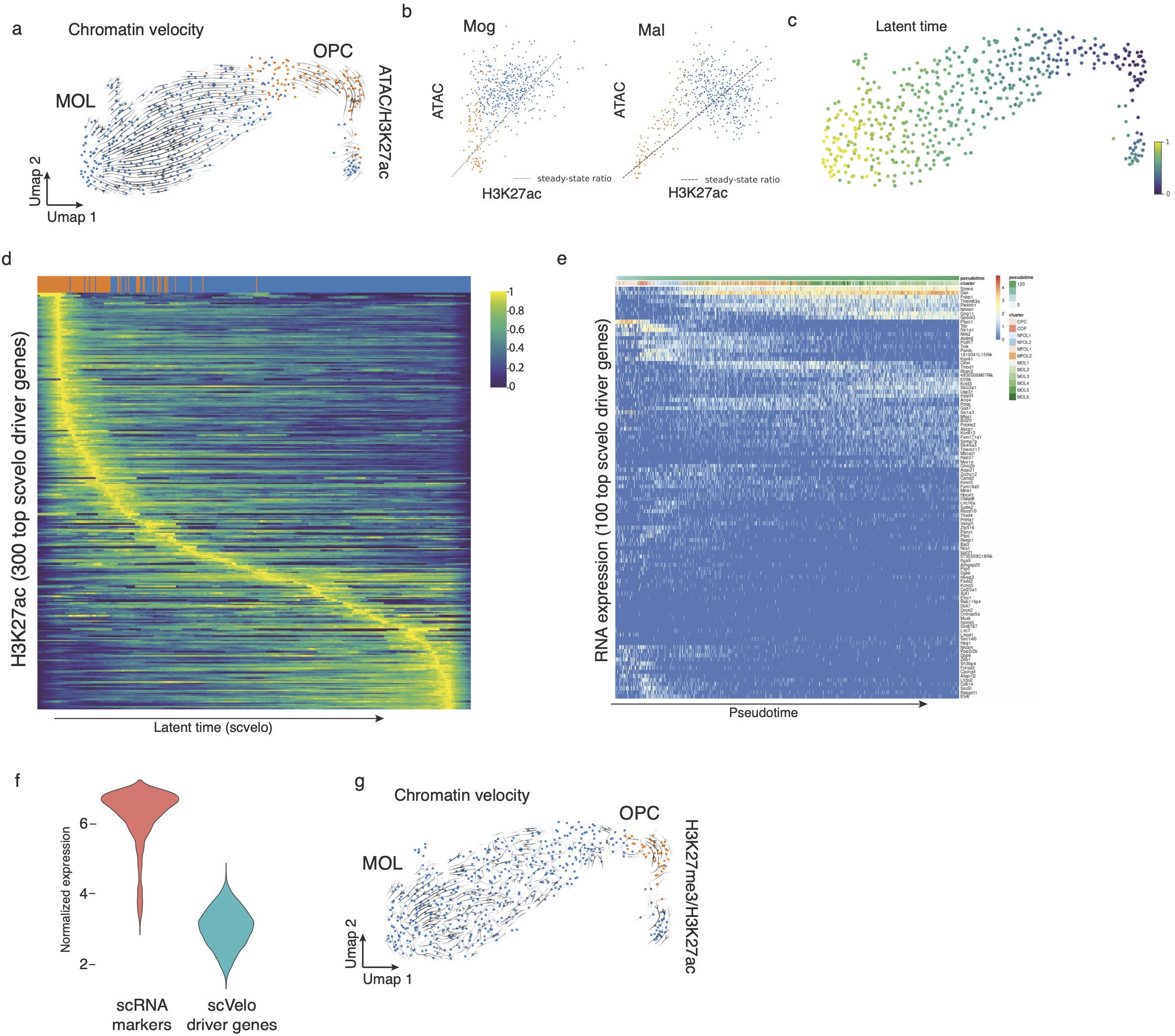
nano-CT based chromatin velocity analysis. **a.** UMAP projection and chromatin velocity visualization. The chromatin velocity was calculated by using ATAC-seq gene by cell matrix as input into the unspliced layer and H3K27ac gene by cell matrix into the spliced layer and then running scvelo algorithm using default parameters. **b**. Phase plots of ATAC-seq and H3K27ac signal for key genes associated with oligodendrocyte differentiation (*Mal, Mag*) **c**. UMAP projection of the latent time calculated by the scvelo algorithm. **d**. Heatmap showing H3K27ac signal normalized with sctransform^35^. Rows depict individual top velocity driver genes, sorted by time of value with maximum intensity and columns depict individual cells sorted by latent time. **e**. Heatmap representing gene expression profiles measured by scRNA-seq^31^ in the oligodendrocytes lineage. Rows depicts individual genes, clustered by similarity and columns depict single cells ordered in pseudotime. **f**. Violin plot showing normalized expression of set of marker genes identified in scRNA-seq dataset, and normalized expression of a set of genes identified as the key driver genes by scvelo. **g**. UMAP projection and velocity vectors projection of chromatin velocity calculated using H3K27ac gene by cell matrix used as input into unspliced layer and H3K27me3 gene by cell matrix used as input into the spliced layer and then running scvelo algorithm using default parameters.

We then identified the key velocity driver genes (Extended data figure 11a, Supplementary table 2) and plotted normalized H3K27ac, ATAC signal and velocity along inferred latent time (Figure 6c) highlighting the dynamic changes in chromatin during oligodendrocytes differentiation (Figure 6d, Extended data figure 11b, c). We validated that the majority of the driver genes also show variable gene expression (measured by scRNA-seq) in the oligodendrocyte lineage^31^ (Figure 6e). Interestingly, the pattern of their induction occurs at different stages during pseudotime (Figure 6d) and these driver genes are relatively lowly expressed, compared to the typical marker genes identified by scRNA-seq (Figure 6f). The GO terms associated with these genes were nervous system development, cell development, neurogenesis or generation of neurons (Extended data figure 11d), suggesting that the set of genes identified through chromatin velocity comprised non-canonical oligodendrocyte differentiation genes. This highlights the strength of multimodal chromatin profiling and chromatin velocity modeling in identification of genes, which are dynamically regulated through changes in the chromatin landscape, but difficult to pick up through gene expression profiling.

Finally, we attempted to model RNA velocity using other combinations of chromatin modalities that do not follow the same causal relationship as ATAC/H3K27ac as input into the velocity analysis. H3K27ac/H3K27me3 are anticorrelated and mutually exclusive histone marks and this combination of modalities did not correctly predict the oligodendrocyte differentiation trajectory (Figure 6g). Thus, anti-correlated active and repressive chromatin marks might require other modelling or data pre-processing strategies to correctly predict future cell states.

## Discussion

Several studies have attempted to profile multiple epigenetic modalities at the same time through direct measurement^21,22^ or imputation from other co-profiled modality^20,23^. Here we describe nano-CT, a new multimodal scCUT&Tag protocol that allows profiling thousands of single cells while 1) significantly reducing the requirements for input material by around 10-fold; 2) increasing the number of fragments detected per cell by 16-fold and 3) allowing the simultaneous probing in each single cell of 3 epigenomic modalities – chromatin accessibility (ATAC), H2K27ac and H3K27me3.

We applied nano-CT to the mouse juvenile brain demonstrated that the simultaneous detection of ATAC, repressive and active histone marks can provide further deconvolution of cell heterogeneity. Moreover, we found that individual modalities are not equally effective in the identification of cell identity and that histone marks can be more informative than for example open chromatin regions, given similar data quality. Finally, we show that nano-CT could for the first time provide direct measurement of time-resolved dynamics of chromatin opening, H3K27ac deposition at the individual enhancer level.

Previously, integration of scCUT&Tag-Pro datasets suggested that H3K27me3 repressive states might be more heterogeneous than what gene expression allows to infer^23^. Here we show that multimodal nano-CT permits deconvolution of H3K27me3 states in a continuous and dynamic biological process such as the oligodendrocyte differentiation. Sequential waves of H3K27me3 occur in pseudotime, at distinct loci regulating distinct modules necessary for efficient OL lineage progression and differentiation.

The use of fusion proteins of Tn5 with single-chain nanobodies recognizing mouse and rabbit antibodies confers a unique versatility to nano-CT. The modular design of the presented nano-Tn5 fusions permit adopting this strategy to virtually any combination of histone marks or chromatin binding proteins. Here we show that the simultaneous, coupled profiling of three epigenetic modalities can provide unique insights into dynamic chromatin remodeling. Tn5 fusion with other single-chain nanobodies might allow to analyze more than three epigenomic modalities in the same cell, at the same time, which is bound to provide further insights into chromatin dynamics. We anticipate that measurement of levels of other histone marks, or chromatin binding factors (transcription factors, chromatin remodelers, RNA polymerase etc.) with nano-CT can provide mechanistic insights into multitude of processes such as activation of enhancers, promoters, initiation of transcription or bivalency.

## Methods

### Animals

The mouse line used in this study was generated by crossing Sox10:Cre animals^7^ (The Jackson Laboratory mouse strain 025807) on a C57BL/6j genetic background with RCE:loxP (EGFP) animals^8^ (The Jackson Laboratory mouse strain 32037-JAX) on a C57BL/6xCD1 mixed genetic background. Females with a hemizygous Cre allele were mated with males lacking the Cre allele, while the reporter allele was kept in hemizygosity or homozygosity in both females and males. In the resulting Sox10:Cre-RCE:LoxP (EGFP) animals the entire OL lineage was labeled with EGFP. Breeding with males containing a hemizygous Cre allele in combination with the reporter allele to non-Cre carrier females resulted in offspring where all cells were labeled with EGFP and was therefore avoided. All animals were free from mouse viral pathogens, ectoparasites and endoparasites and mouse bacteria pathogens. Mice were kept with the following light/dark cycle: dawn 6:00-7:00, daylight 7:00-18:00, dusk 18:00-19:00, night 19:00-6:00 and housed to a maximum number of 5 per cage in individually ventilated cages (IVC sealsafe GM500, tecniplast). Cages contained hardwood bedding (TAPVEI, Estonia), nesting material, shredded paper, gnawing sticks and card box shelter (Scanbur). The mice received regular chow diet (either R70 diet or R34, Lantmännen Lantbruk, Sweden). Water was provided by using a water bottle, which was changed weekly. Cages were changed every other week. All cage changes were done in a laminar air-flow cabinet. Facility personnel wore dedicated scrubs, socks and shoes. Respiratory masks were used when working outside of the laminar air-flow cabinet. Animals were sacrificed at juvenile stages (P19) and both sexes were included in the study. All experimental procedures on animals were performed following the European directive 2010/63/EU, local Swedish directive L150/SJVFS/2019:9, Saknr L150 and Karolinska Institutet complementary guidelines for procurement and use of laboratory animals, Dnr. 1937/03-640. The procedures described were approved by the local committee for ethical experiments on laboratory animals in Sweden (Stockholms Norra Djurförsöksetiska nämnd), lic. nr. 144/16 and 1995_2019.

### Antibodies

The following antibodies were used in the multimodal nano-CT experiments: mouse anti-H3K27me3 (Abcam, Ab6002), rabbit anti-H3K27ac (Abcam Ab177178). Anti-H3K27me3 (Cell Signaling, 9733T) was used in single modality nano-CT experiment.

### Nanobody-Tn5 fusion design and production

The sequence of secondary nanobodies was taken from Pleiner et al.^24^ and fused to the sequence of hyperactive Tn5 transposase. For anti-rabbit-Tn5 fusion with nanobody TP897 and for anti-mouse-Tn5 fusion with nanobody TP1170 was designed. The full sequence of the fusion proteins can be found in supplementary materials (nanobody_Tn5_sequences.fa). The fulllength fusion protein sequences were ordered from Twist Bioscience cloned into twist expression vector. The constructs were transformed into BL21 (DE3) Star *E. coli*, inoculated into an overnight culture and grown in TB medium supplemented with 8g/l glycerol at 30°C with 175 RPM shaking in the presence of 0.4% glucose. The next day the cultures were grown in the LEX system with start in the morning starting with cultivation at 37°C. The OD was measured at different times and the temperature was switched to 18°C, when the culture reached OD=2. The protein expression was induced at approximately OD 3 (IPTG, final concentration 0.5 mM). Protein expression continued overnight before the cells were harvested by centrifugation (10 min at 4500 × g). IMAC lysis buffer (1.5 ml buffer per gram cell pellet) and the complete stock solution (1 tablet Complete EDTA-free (protease inhibitor cocktail, Roche) and 50 μl benzonase nuclease cell resuspension (PSF) per 1 ml; 1 ml per 1.5 l culture) were added and the cell pellets were re-suspended on a shaker table (cold room). The resuspended cell pellets were stored at −80°C.

The frozen cell pellets were thawed, extra benzonase was added and the cells were disrupted by pulsed sonication (4s/4s 4 min, 80% amplitude). The sonicated lysates were centrifuged (20 min at 49000 × g) and the soluble fractions were decanted and filtered through 0.45 μm filters. The samples were loaded onto the ÄKTA Xpress and purified overnight.

Buffers: Lysis buffer: 100 mM HEPES, 500 mM NaCl, 10% glycerol, 10 mM imidazole, 0.1 mM EDTA, 0.1% Triton X-100, 0.5 mM TCEP, pH 8.0; Wash 1 buffer: 20 mM HEPES, 500 mM NaCl, 10% glycerol, 10 mM imidazole, 0.1 mM EDTA, 0.1% Triton X-100, 0.5 mM TCEP, pH 7.2; Wash 2 buffer: 20 mM HEPES, 1000 mM NaCl, 10% glycerol, 50 mM imidazole, 0.1 mM EDTA, 0.1% Triton X-100, 0.5 mM TCEP, pH 7.2; Elution buffer: 20 mM HEPES, 500 mM NaCl, 10% glycerol, 500 mM imidazole, 0.1 mM EDTA, 0.1% Triton X-100, 0.5 mM TCEP, pH 7.2; Gel filtration buffer: 50 mM HEPES, 500 mM NaCl, 10% glycerol, 0.1 mM EDTA, 0.1% Triton X-100, 1 mM DTT, pH 7.2; Final batch buffer: 50 mM HEPES, 500 mM NaCl, 50% glycerol, 0.1 mM EDTA, 0.1% Triton X-100, 1 mM DTT, pH 7.2; IMAC Column: 5 ml HisTrap HP (GE Healthcare); Gel Filtration Column: HiLoad 16/60 Superdex 200 (GE Healthcare). Selected fractions were examined on SDS-PAGE gels before pooling. Fractions containing the target proteins were pooled and concentrated with Vivaspin concentration filters (Vivascience, 50 kDa cut off) at 4000 × g. The samples were then diluted with buffer containing 60% glycerol to get 50% in the final buffer. The final concentration was measured by Abs 280 nm (Nanodrop), the protein was flash frozen in aliquots á 200 μL in liquid nitrogen and stored at −80°C.

### Tn5 loading

Nanobody-Tn5 fusion protein was loaded with barcoded oligonucleotides. First, equimolar mixture of 100 uM forward oligos (Tn5_P5_MeA_BcdX_0N, Tn5_P5_MeA_BcdX_1N, Tn5_P5_MeA_BcdX_2N, Tn5_P5_MeA_BcdX_3N; supplementary table 1) was mixed (5 ul each) and with equimolar amount of 100 uM of Tn5_Rev oligo (20ul). The oligo mixture was denatured for 5 minutes at 95°C and annealed by slowly ramping down the temperature at 0.1°C per second in a thermocycler. The nano-Tn5 was loaded by mixing the following: 8ul of annealed (P5) oligonucleotides, 42 ul of Glycerol, 44.1 ul of 2× dialysis buffer (100 mM HEPES-KOH pH7.2; 200 mM NaCl; 0.2 mM EDTA; 2 mM DTT (add fresh); 0.2% Triton-X; 20% Glycerol) 5.9 ul anti-mouse-nano-Tn5 (5mg/ml, 67.6 uM) or 8ul of annealed oligonucleotides, 42 ul of Glycerol, 45.7 ul of 2x dialysis buffer (100 mM HEPES-KOH pH7.2; 200 mM NaCl; 0.2 mM EDTA; 2 mM DTT; 0.2% Triton-X; 20% Glycerol) 4.3 ul anti-rabbit-nano-Tn5 (6.8 mg/ml, 93uM) to get final 2uM loaded nano-Tn5 dimer.

Non-fused Tn5 (as in ATAC-seq) was loaded using Tn5_P7_MeB standard oligonucleotides (referred to as P7 Tn5). First the 100 uM Tn5_P7_MeB oligo was mixed with equimolar amount of 100 uM Tn5_Rev. The oligo mixture was denatured for 5 minutes at 95°C and annealed by slowly ramping down the temperature at 0.1°C per second in a thermocycler. The Tn5 was loaded by mixing the following: 8ul of annealed oligonucleotides (P7), 43.12 ul of Glycerol, 42.6 ul of 2x dialysis buffer (100 mM HEPES-KOH pH7.2; 200 mM NaCl; 0.2 mM EDTA; 2 mM DTT (add fresh); 0.2% Triton-X; 20% Glycerol) 6.28 ul Tn5 (3.5mg/ml, 63.64 uM).

### Tissue dissociation

Mouse were sacrificed, perfused with aCSF buffer (87 mM NaCl, 2.5 mM KCl,1.25 mM NaH2PO4, 26 mM NaHCO_3_, 75 mM Sucrose, 20 mM Glucose, 0.5 mM CaCl_2_ and 4 mM MgSO_4_) and brain was removed. The brain was dissociated into single cell suspension using the Neural Tissue Dissociation Kit P (Miltenyi Biotec, 130-092-628) according to the manufacturers protocol. For mouse older than P7, myelin was removed using debris removal solution (Miltenyi Biotec, 130-109-398) according to manufacturer’s instruction. Single cell suspension was filtered through 50 um cell strainer.

### nano-CT

12 500 – 250 000 cells were counted, centrifuged for 5 minutes at 500x g, resuspended in 200 ul of antibody buffer (20 mM HEPES pH 7.5, 150 mM NaCl, 2mM EDTA, 0.5 mM Spermidine, 0.05% Digitonin, 0.01 % NP-40, 1x Protease inhibitors, 2% BSA) and incubated for 3 minutes on ice to extract nuclei. Nuclei were then centrifuged at 600x g for 3minutes and resuspended in 100ul of antibody buffer pre-mixed with 1:100 diluted primary mouse antibody, 1:100 diluted primary rabbit antibody, 1:100 diluted anti-rabbit-nano-Tn5 loaded with barcoded P5 oligonucleotide and 1:100 diluted anti-mouse-nano-Tn5 loaded with a different barcoded P5 oligonucleotide. The barcodes that were used were recorded. The sample was then incubated at 4°C on a roller overnight.

After overnight incubation, the nuclei were centrifuged at 600× g for 3 minutes and washed twice with Dig-300 buffer (20 mM HEPES pH 7.5, 300 mM NaCl, 0.5 mM Spermidine, 0.05% Digitonin, 0.01 % NP-40, 2% BSA, 1× Protease inhibitors). After that, the nuclei were resuspended by pipetting in 200ul of tagmentation buffer (20 mM HEPES pH 7.5, 300 mM NaCl, 0.5 mM Spermidine, 0.05% Digitonin, 0.01 % NP-40, 2%BSA, 1× Protease inhibitors, 10 mM MgCl_2_) and incubated for 1 hour at 37°C. The sample was mixed by pipetting after 30 minutes of the incubation to prevent sedimentation. The reaction was stopped by addition of 200 ul of 1× Diluted nuclei buffer (Chromium Next GEM Single Cell ATAC Library & Gel Bead Kit v1.1; 10× Genomics) supplemented with 2% BSA (1x DNB/BSA) and 12.5 mM EDTA. The nuclei were sedimented by centrifugation at 600× g for 3 minutes and washed twice with 200ul of 1× DNB supplemented with 2%BSA. After the second wash and centrifugation, around 180ul of the 1x DNB/BSA supernatant was removed from the sample. The nuclei were resuspended in the remaining 20 ul of 1xDNB/BSA, 2ul were mixed with 8 ul of trypan blue and counted manually using a counting chamber. 16,000 nuclei were used to load the 10x chromium chip.

### nano-CT with ATAC-seq

ATAC-seq was performed as described in^32^ (Omni-ATAC) with minor modifications. Tn5 was loaded with barcoded MeA/Me-Rev oligonucleotides omitting the MeB/MeR (see part Tn5 loading and supplementary table 1). The number of cells used as input was scaled up to 200,000. The cells were counted, centrifuged for 5 minutes at 500× g, resuspended in 200ul of ATAC resuspension buffer (ARB, 10mM Tris pH 7.5, 10mM NaCl, 3mM MgCl2 supplemented with 0.1% NP-40, 0.1% Tween-20 and 0.01% Digitonin), pipetted up and down 3 times and incubated on ice for 3 minutes. Then 1 ml of ATAC wash buffer (ARB, supplemented with 0.1 Tween-20) was added, and mixed by inverting the tube 3 times. Nuclei were pelleted by centrifugation at 500× g for 10 minutes at 4°C on swinging bucket rotor centrifuge with appropriate adaptors for microtubes. Supernatant was aspirated and cell pellet as resuspended in 200 ul of transposition mix (100 ul 1× TD buffer, 66ul 1x PBS, 2ul 10% Tween-20, 2ul 1% digitonin, 10 ul Tn5 transposase loaded with barcoded MeA/Rev (P5) oligonucleotides, 20ul water), pipetted up and down 5 times gently to mix and incubated at 37°C for 30 minutes with 800 rpm mixing. After the tagmentation was finished, the nuclei were centrifuged at 500x g for 10 minutes, supernatant was removed, and nuclei were washed with 200ul of CT antibody buffer (20 mM HEPES pH 7.5, 150 mM NaCl, 2mM EDTA, 0.5 mM Spermidine, 0.05% Digitonin, 0.01 % NP-40, 1× Protease inhibitors, 2% BSA). Then, the nuclei were pelleted by centrifugation for 3 minutes at 600× g, the washed again with 200 ul of antibody buffer and centrifuged for 3 minutes at 600× g. The pellet was then resuspended in 100ul of antibody buffer pre-mixed with 1:100 diluted primary mouse antibody, 1:100 diluted primary rabbit antibody, 1:100 diluted anti-rabbit-nano-Tn5 loaded with barcoded P5 oligonucleotide and 1:100 diluted anti-mouse-nano-Tn5 loaded with a different barcoded P5 oligonucleotide. The barcodes that were used were recorded. The sample was then incubated at 4°C on a roller overnight.

After overnight incubation, the nuclei were centrifuged at 600x g for 3 minutes and washed twice with Dig-300 buffer (20 mM HEPES pH 7.5, 300 mM NaCl, 0.5 mM Spermidine, 0.05% Digitonin, 0.01 % NP-40, 2% BSA, 1x Protease inhibitors). After that, the nuclei were resuspended by pipetting in 200ul of tagmentation buffer (20 mM HEPES pH 7.5, 300 mM NaCl, 0.5 mM Spermidine, 0.05% Digitonin, 0.01 % NP-40, 2%BSA, 1× Protease inhibitors, 10 mM MgCl2) and incubated for 1 hour at 37°C. The sample was mixed by pipetting after 30 minutes of the incubation to prevent sedimentation. The reaction was stopped by addition of 200 ul of 1× Diluted nuclei buffer (Chromium Next GEM Single Cell ATAC Library & Gel Bead Kit v1.1; 10x Genomics) supplemented with 2% BSA (1× DNB/BSA) and 12.5 mM EDTA. The nuclei were sedimented by centrifugation at 600× g for 3 minutes and washed twice with 200ul of 1× DNB supplemented with 2%BSA. After the second wash and centrifugation, around 180ul of the 1× DNB/BSA supernatant was removed from the sample. The nuclei were resuspended in the remaining 20 ul of 1xDNB/BSA, 2ul were mixed with 8 ul of trypan blue and counted manually using a counting chamber. 16,000 nuclei were used to load the 10× chromium chip.

### Single cell indexing and library prep

Single cell indexing was performed using Chromium Next GEM Single Cell ATAC Library & Gel Bead Kit v1.1 (10x Genomics). Up to 8 ul of Nuclei suspension (filled up to 8 ul with 1× DNB/BSA) was mixed with 7ul of 10× ATAC buffer B, 56.5 ul of barcoding reagent B, 1.5 ul of reducing agent B and 1ul of barcoding enzyme. 70ul of the mastermix was loaded on Chromium Next GEM Chip H together with 50 ul of gel beads and 40 ul of partitioning oil according to manufacturer’s instructions followed by single cell partitioning using the chromium controller and GEM incubation according to manufacturer’s instructions (Chromium Next GEM Single Cell ATAC Reagent Kits v1.1 (Steps 2.0-2.5)).

Upon completion, the sample was recovered by post-GEM incubation cleanup with Dynabeads MyOne SILANE and SPRIselect beads according to Chromium Next GEM Single Cell ATAC Reagent Kits v1.1 (Steps 3.0-3.2)

After elution from SPRIselect beads, 40ul of sample was recovered and 2ul were used to measure the concentration using Qubit dsDNA HS Assay kit (ThermoFisher). The rest of the sample was used for P7 tagmentation by mixing the following reaction: 38ul template DNA, 50 ul 2× TD buffer (20 mM Tris pH 7.5, 10mM MgCl2, 20% Dimethylformamide), 1 ul of MeB-only loaded standard Tn5 (P7 Tn5; Typicaly 0.5-1 ul of P7 Tn5 in single-cell experiment; see Tn5 loading section for details on loading conditions) per 10ng of template DNA and dH2O up to 100 ul and incubation for 30 minutes at 37°C in a thermocycler. After that, the sample was purified with DNA Clean and concentrator-5 (Zymo) using 500 ul of binding buffer and eluted in 40 ul of Zymo elution buffer. The purified DNA was used as a template for the final PCR amplification according to Chromium Next GEM Single Cell ATAC Reagent Kits v1.1 (Step 4.1) (40 ul template, 50 ul AMP mix, 7.5 ul SI-PCR primer B and 2.5 ul of individual Single index N Set A primer) and amplified using standard 10× scATAC-seq PCR amplification program (1. 98°C 45s; 2. 98°C 20s; 3. 67°C 30s; 4. 72°C 20s N; 5. 72°C 1min; 6. 4°C hold) and amplified for 11-15 cycles. The amplified product was purified with SPRIselect reagent (Beckman-Coulter) using 0.4× and 1.2× two-sided purification according to the Chromium Next GEM Single Cell ATAC Reagent Kits v1.1 (Step 4.2). The size distribution and concentration of the library was assessed using Qubit dsDNA HS Assay kit (ThermoFisher) and bioanalyzer high sensitivity DNA kit.

### Sequencing setup

The libraries were sequenced using custom read1 (R1_seq) and index2 (I2_seq) primers (Supplementary Table1) with custom read lengths 36-8-48-36 (R1-I1-I2-R2).

### Data analysis

#### Preprocessing

Raw fastq files were demultiplexed into individual fastq files for each modality using debarcode.py script with 1 allowed mismatch in barcode. Fastq files were then used as input into cellranger 1.2.0 pipeline, with modifications. Peaks were called using pseudobulk bam files outputted from cellranger (outs/possorted_bam.bam) using MACS2 with parameters --keep-dup=1 --llocal 100000 --min-length 1000 --max-gap 1000 --broad-cutoff=0.1. Custom script was used to identify single cells based on number of reads per cell and fraction of reads in peaks (counted using bedtools intersect) (Extended data figure 3). Seurat object was then created for each modality and replicate separately using R function FeatureMatrix and peaks called previously. Seurat objects were then merged per modality (ATAC, H3K27ac, H3K27me3) and across the modalities.

#### Dimensionality reduction and clustering

All dimensionality reduction and clustering was performed using package Seurat v4.0.5 and Signac v1.4.0 under R version 4.1.1. Dimensionality reduction was performed using latent semantic indexing (LSI) and uniform manifold approximation and projection (UMAP). Clusters were annotated through integration with scRNA-seq data (mouse adolescent brain atlas)^33^ and manual annotation based on marker genes. Bigwig files per cell cluster were constructing by filtering the possorted_bam.bam file using single-cell barcodes by custrom script creating bam file per cluster and bamCoverage script from deeptools.

#### Downstream analysis

Metagene plots were generated using deeptools package v3.5.1 using computeMatrix and plotHeatmap scripts. Scores for individual peak regions were generated using deeptools multiBigwigSummary script. GO analysis was performed using R package clusterProfiler v4.2.2.

#### Chromatin dynamics analysis

Pseudotime in the oligodendrocyte lineage was calculated using R package slingshot v2.2.0^34^. Chromatin velocity was performed using python package scvelo v0.2.4 by directly using ATAC-seq and H3K27ac matrix gene by cell matrix into unspliced and spliced layers respectively. Scvelo analysis was performed using default parameters.

#### Material availability

The plasmids used for purification of nanobody Tn5-fusions have been submitted to Addgene (#183637, #183638).

#### Code availability

Custom code, scripts and analysis pipeline can be found at https://github.com/mardzix/bcd_nano_CUTnTag.

## Supporting information

Supplementary table 1. List of oligonucleotides used in this study

Supplementary table 2. List of scvelo driver genes in the oligodendrocyte lineage

Protein sequences of nanobody-Tn5 fusion proteins used in this study

## Data availability

Raw data was deposited in GEO under accession GSE198467. The following publicly available datasets were used in this study <u>GSE75330</u> (scRNA-seq of oligodendrocyte lineage), <u>GSE163532 (scCUT&Tag in the juvenile mouse brain)</u>, https://storage.googleapis.com/linnarsson-lab-loom/l5_all.loom (scRNA-seq, Mouse juvenile brain atlas),

## Acknowledgements

We would like to thank Tony Jimenez-Beristain for writing laboratory animal ethics permit 1995_2019 and assistance with animal experiments, Leslie Kirby, Mandy Meijer, Chao Zheng and Petra Kukanja for assistance with animal experiments, Tomas Nyman, Emilia Strandback, Henry Ampah-Korsah and Karolinska Institute Protein Science facility for the nanobody-Tn5 constructs and the protein purification, Simon Elsässer for critical discussions, Eneritz Agirre, Mukund Kabbe and Neemat Mahmud for proofreading and input on the manuscript, and the staff at Comparative Medicine-Biomedicum. The authors acknowledge support from the National Genomics Infrastructure in Stockholm funded by Science for Life Laboratory, the Knut and Alice Wallenberg Foundation and the Swedish Research Council, and Swedish National Infrastructure for Computing /Uppsala Multidisciplinary Center for Advanced Computational Science for assistance with massively parallel sequencing and access to the UPPMAX computational infrastructure. M.B. is funded by the Vinnova Seal of Excellence Marie-Sklodowska Curie Actions grant RNA-centric view on Oligodendrocyte lineage development (RODent). Work in G.C.-B.’s research group was supported by the Swedish Research Council (grant 2015-03558 and 2019-01360), the European Union (Horizon 2020 Research and Innovation Programme/European Research Council Consolidator Grant EPIScOPE, grant agreement number 681893), the Swedish Brain Foundation (FO2017-0075), the Swedish Cancer Society (Cancerfonden; 190394 Pj), Knut and Alice Wallenberg Foundation (grant 2019-0107 and 2019-0089), The Swedish Society for Medical Research (SSMF, grant JUB2019), Göran Gustafsson foundation, Ming Wai Lau Centre for Reparative Medicine and Karolinska Institutet. Some figures were created with BioRender.com.

## Author Contributions

MB and GCB conceived the study, designed the experiments and analysis and wrote the manuscript. MB prepared the figures. MB optimized and performed the nanoCUT&Tag experiments and analyzed the data. All authors contributed and approved the manuscript.

---

**Extended data figure 1. a.** qPCR amplification curve for libraries generated using the same amount of input material using scCUT&Tag method, scCUT&Tag combined with linear preamplification and nano-CT protocol with two-step tagmentation. **b**. Bioanalyser plot showing the size distribution of libraries generated using scCUT&Tag and nano-CT. Typical nucleosome profile can only be seen in the scCUT&Tag library. **c**. UMAP co-embedding of the scCUT&Tag and nano-CT dataset (single modality). The raw count matrices were merged and dimensionality reduction was done on both datasets together. **d**. Violin plot showing fraction of reads in peak regions (FrIP) for data collected by scCUT&Tag and nano-CT.

**Extended data figure 2. a.** Heatmap showing the cell by marker matrix for scCUT&Tag dataset. Top markers were selected based on p-value. **b**. Heatmap showing the cell by marker matrix for nano-CT dataset. Top markers were selected based on p-value. **c**. Boxplot depicting fraction of cells with any signal (counts >1) in the most significantly enriched marker regions for the respective cluster.

**Extended data figure 3.**. Scatter plot showing number of reads per cell (x-axis) versus fraction of reads in peak regions (y-axis). Gaussian mixed model clustering was used to identify clusters of cells and cluster with the highest median of number of reads was selected as valid cells. Valid cells are labeled with cyan, while cells that do not pass QC are red. Contour lines depict density of points withing regions.

**Extended data figure 4. a.** Upset plots showing the overlap between cells that pass QC within the different modalities for combinations of H3K27ac and H3K27me3, and also ATAC, H3K27ac and H3K27me3. **b**. Violin plot showing the distribution of number of reads per cell in nano-CT and scCUT&Tag. **c**. Violin plot showing the distribution of number of reads per in peak regions (FrIP) in nano-CT and scCUT&Tag.

**Extended data figure 5. a.** UMAP co-embedding of CCA-integrated H3K27ac nano-CT together with scRNA-seq dataset. Gene body and promoters were used as gene activity scores for the integration. **b**. UMAP co-embedding of CCA-integrated nano-CT ATAC-seq together with scRNA-seq dataset. Gene body and promoters were used as gene activity scores for the integration.

**Extended data figure 6. a.** UMAP embedding and visualization of marker peak activities for H3K27me3 nano-CT. Exact genomic region together with the closest gene is shown in the plot title. **b**. Violin plot visualization the of peak scores for the same peak regions as in a)

**Extended data figure 7.** Genome browser tracks for all modalities profiled by multimodal nano-CT (ATAC, H3K27ac, H3K27me3) around the **a**. HoxB, **b**. HoxC and **c**. HoxD loci.

**Extended data figure 8.** Confusion matrix of broad cell identities between **a.** ATAC and H3K27ac **b**. ATAC and H3K27me3 and **c**. H3K27ac and H3K27me3. Confusion matrices for fine cluster identities across the different modalities for **d**. ATAC and H3K27ac **e**. ATAC and H3K27me3 and **e**. H3K27ac and H3K27me3.

**Extended data figure 9. a.** Alluvial diagram of corresponding cell identities between the ATAC, H3K27ac and H3K27me3 modalities. **b,c**. UMAP projection and visualization of ATAC, H3K27ac and H3K27me3 signal intensity in single cells at the *Lhx2* **(b)** and Foxb1 **(c)** locus. Grey lines connect the cells with same single-cell barcodes across the different modalities. Clusters telencephalon astrocytes (AST_TE) and non-telencephalon astrocytes (AST_NT) were selected for the visualization.

**Extended data figure 10 a.** UMAP embedding and visualization of pseudotime determined based on the WNN dimensionality reduction. Pseudotime is projected on UMAPs generated by dimensionality reduction of the individual modalities. **b**. Scatter plot depicting meta-region score for all modalities (y-axis) and pseudotime (x-axis). The score was calculated as a sum of normalized score of each individual region. The regions were selected based on p-value (p<0.05, Wilcoxon test) and log fold change > 0 at the marker regions of the H3K27ac modality, and top 200 regions were used. **c**. UMAP embedding of the scRNA-seq data from the mouse brain atlas^33^. 20,000 single cells were sampled randomly from the dataset. Wave 1 and wave2 scores were calculated as a mean of scaled gene expression data for genes identified in figure 5c. **d**. Gene ontology enrichment analysis of genes, which are repressed in the wave 2 of H3K27me3-mediated repression.

**Extended data figure 11 a.** Phase plots of H3K27ac and ATAC, velocity and expression signal for top 10 most likely velocity driver genes identified by scvelo. **b**. Heatmap showing ATAC-seq signal normalized with sctransform^35^. Rows depict individual top velocity driver genes, sorted by time of value with maximum intensity and columns depict individual cells sorted by latent time. **c**. Heatmap showing the chromatin velocity. Rows depict individual top velocity driver genes, sorted by time of value with maximum intensity and columns depict individual cells sorted by latent time. **d**. Gene ontology enrichment analysis of the most important velocity driver genes.

