## Extended data figures 1-11 for "Multimodal chromatin profiling using nanobody-based single-cell CUT&Tag"

### Extended data figure 1

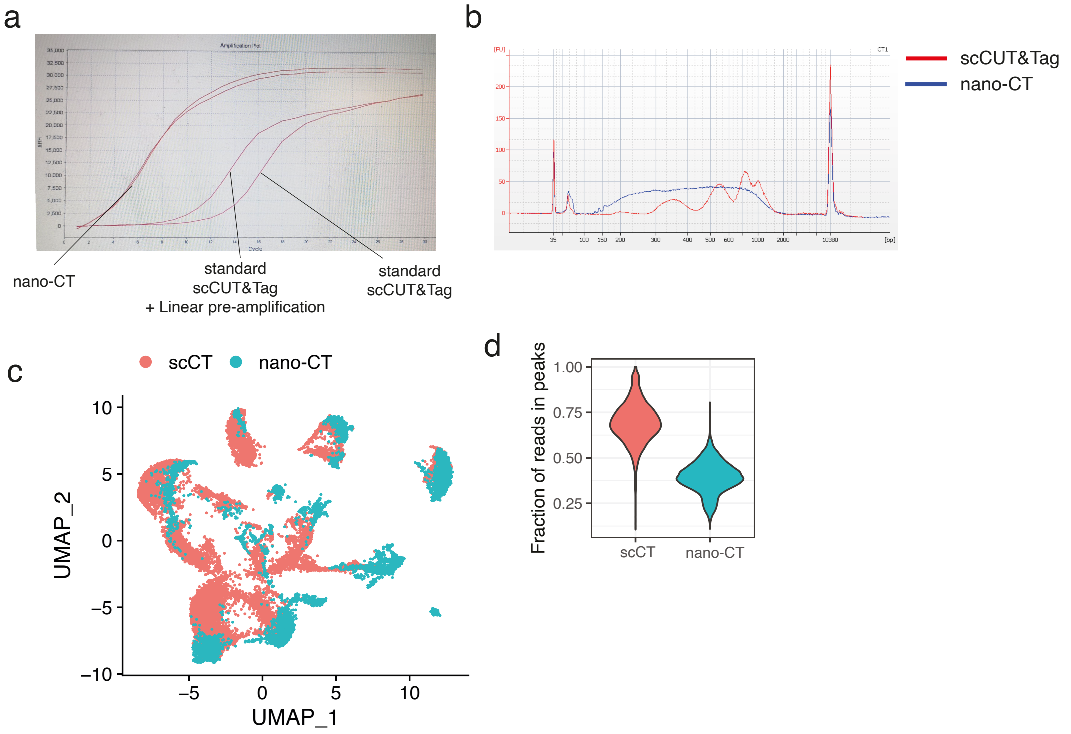

Extended data figure 2

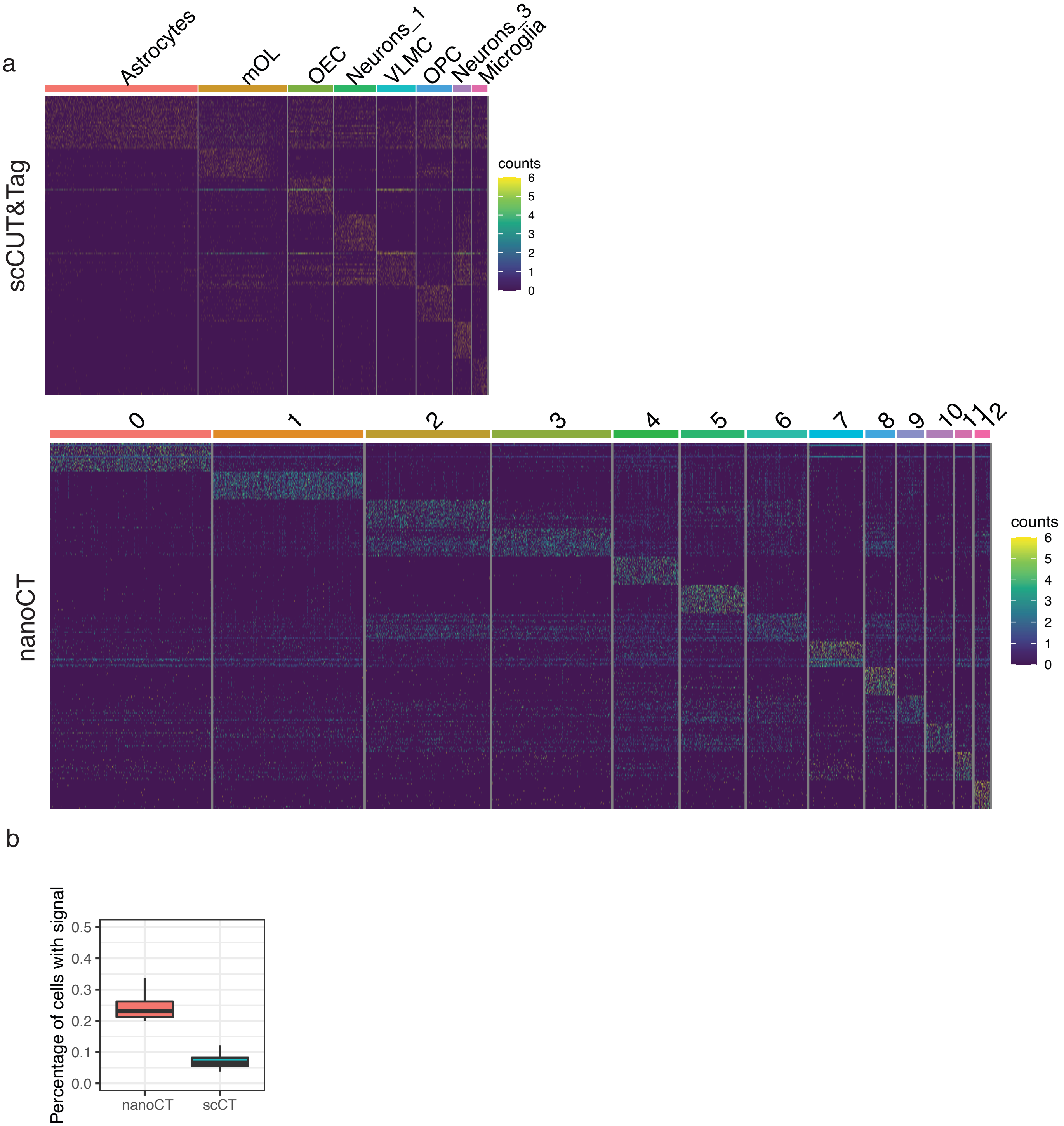

### Extended data figure 3

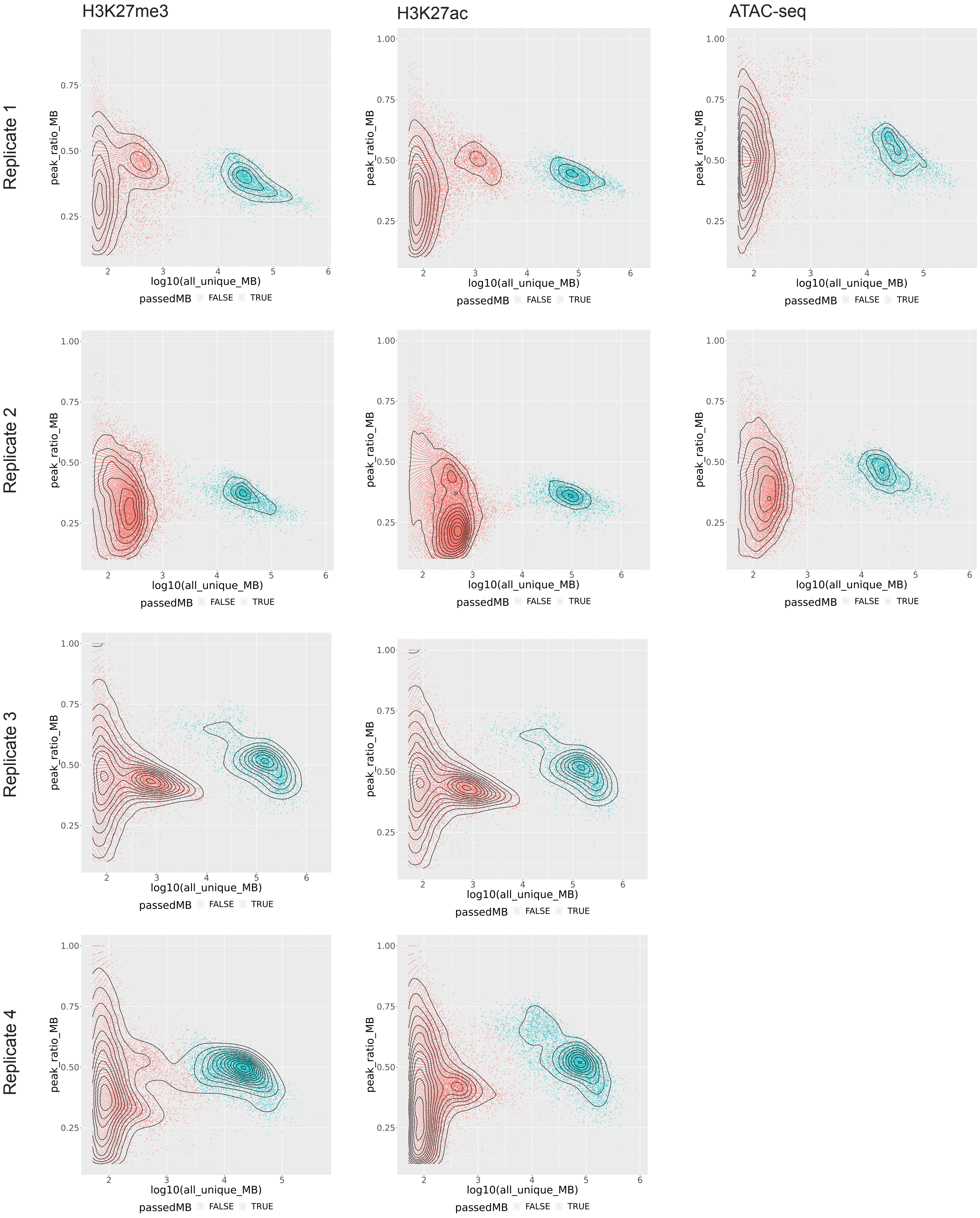

### Extended data figure 4

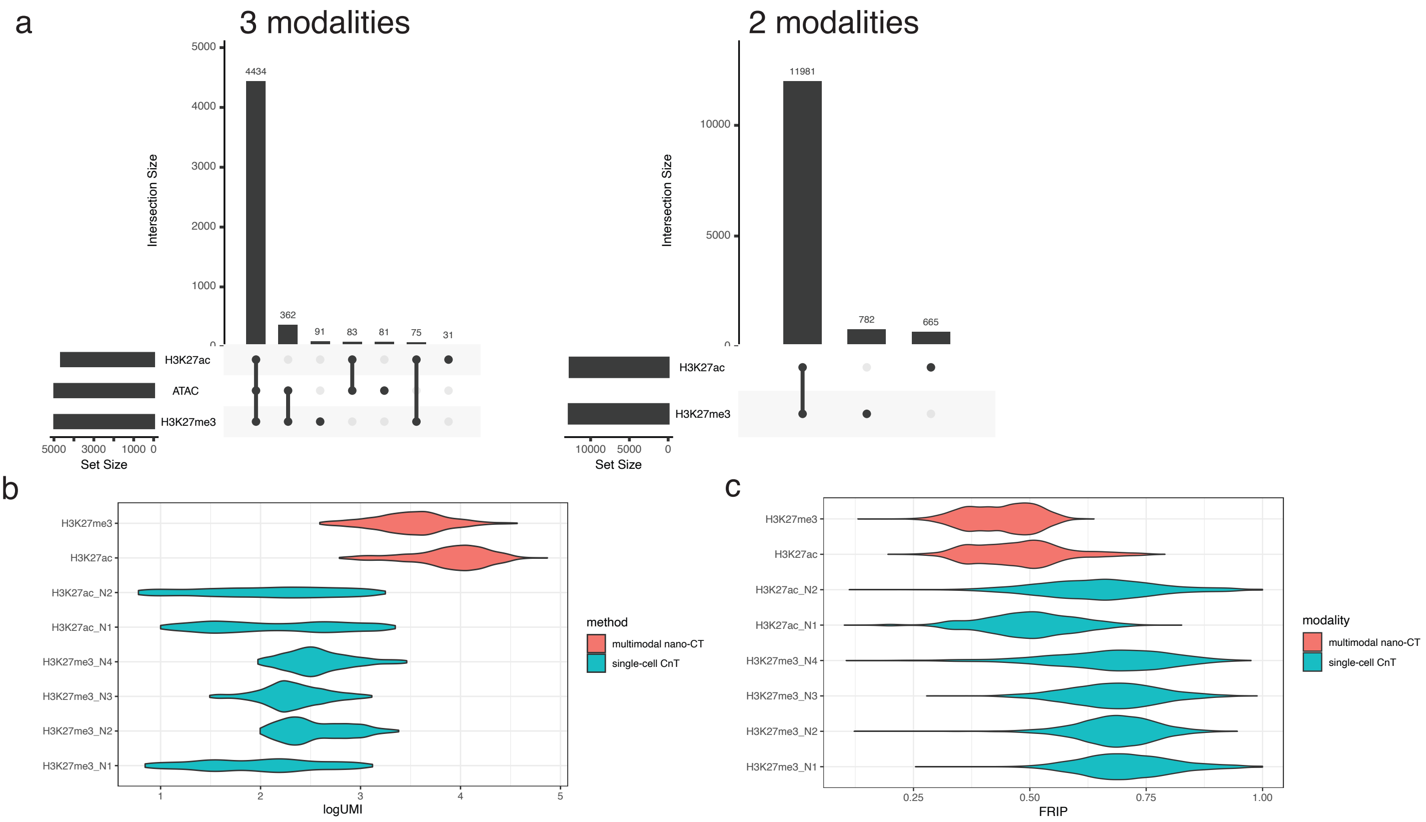

### Extended data figure 5

a

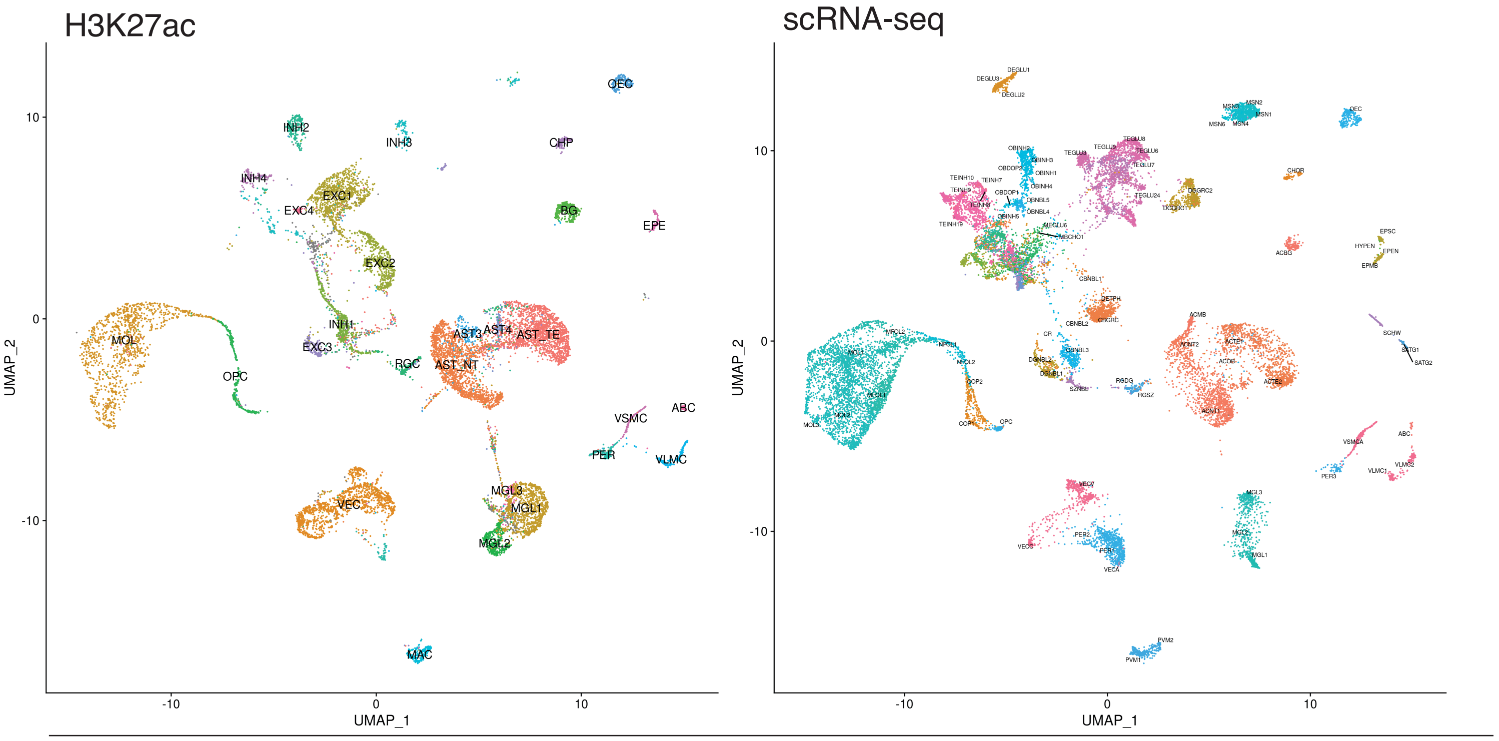


Co-embedding

b

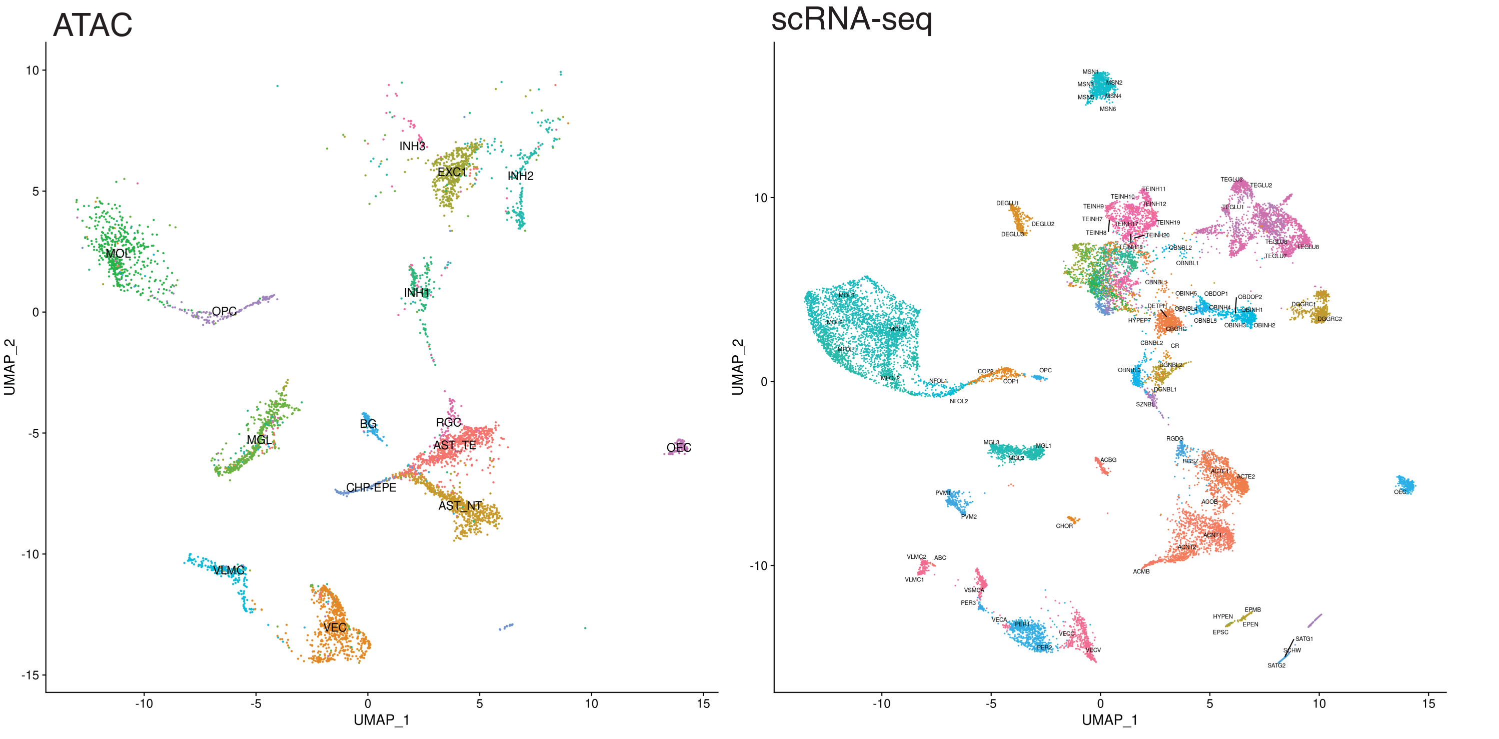


Co-embedding

### Extended data figure 6

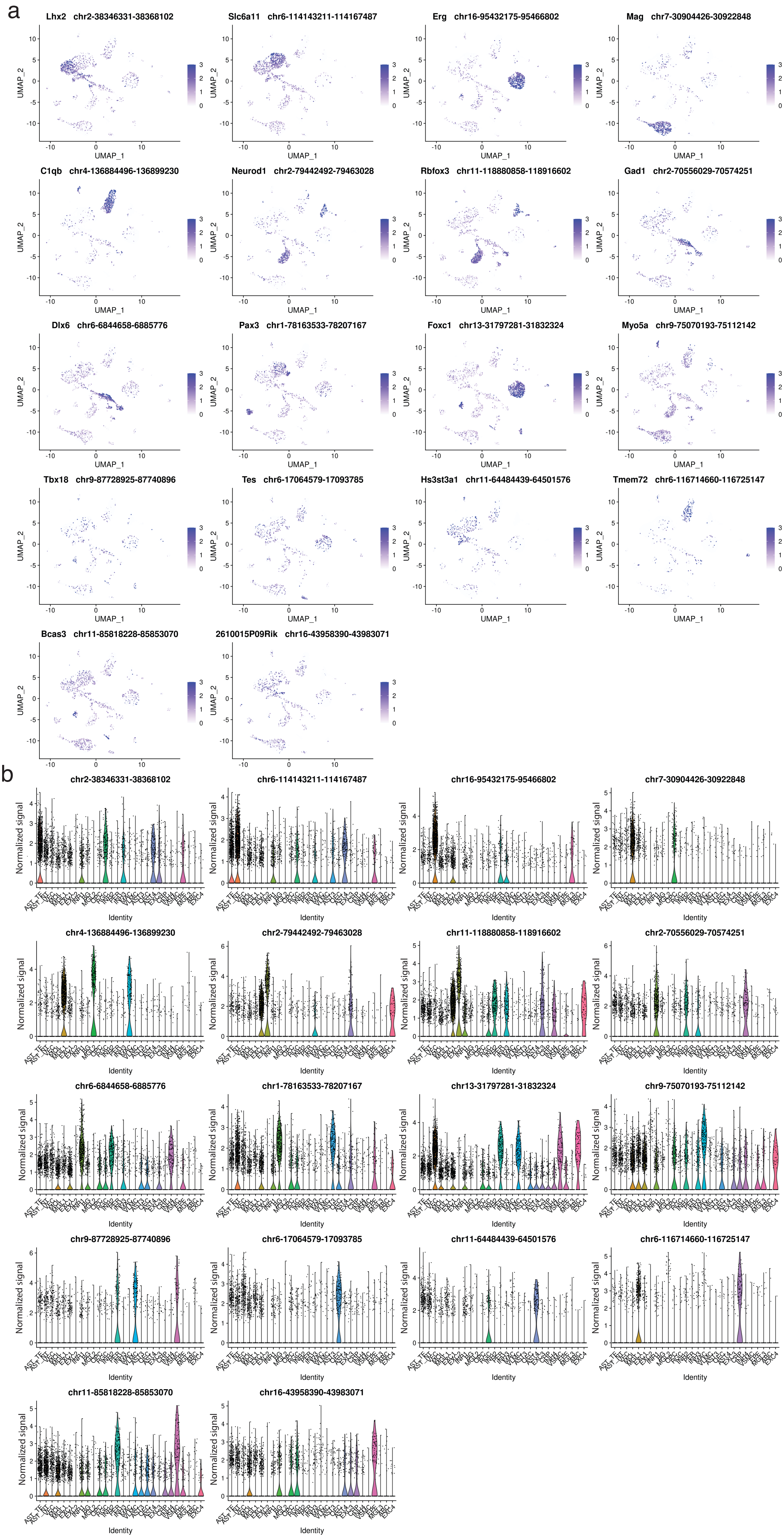

### Extended data figure 7

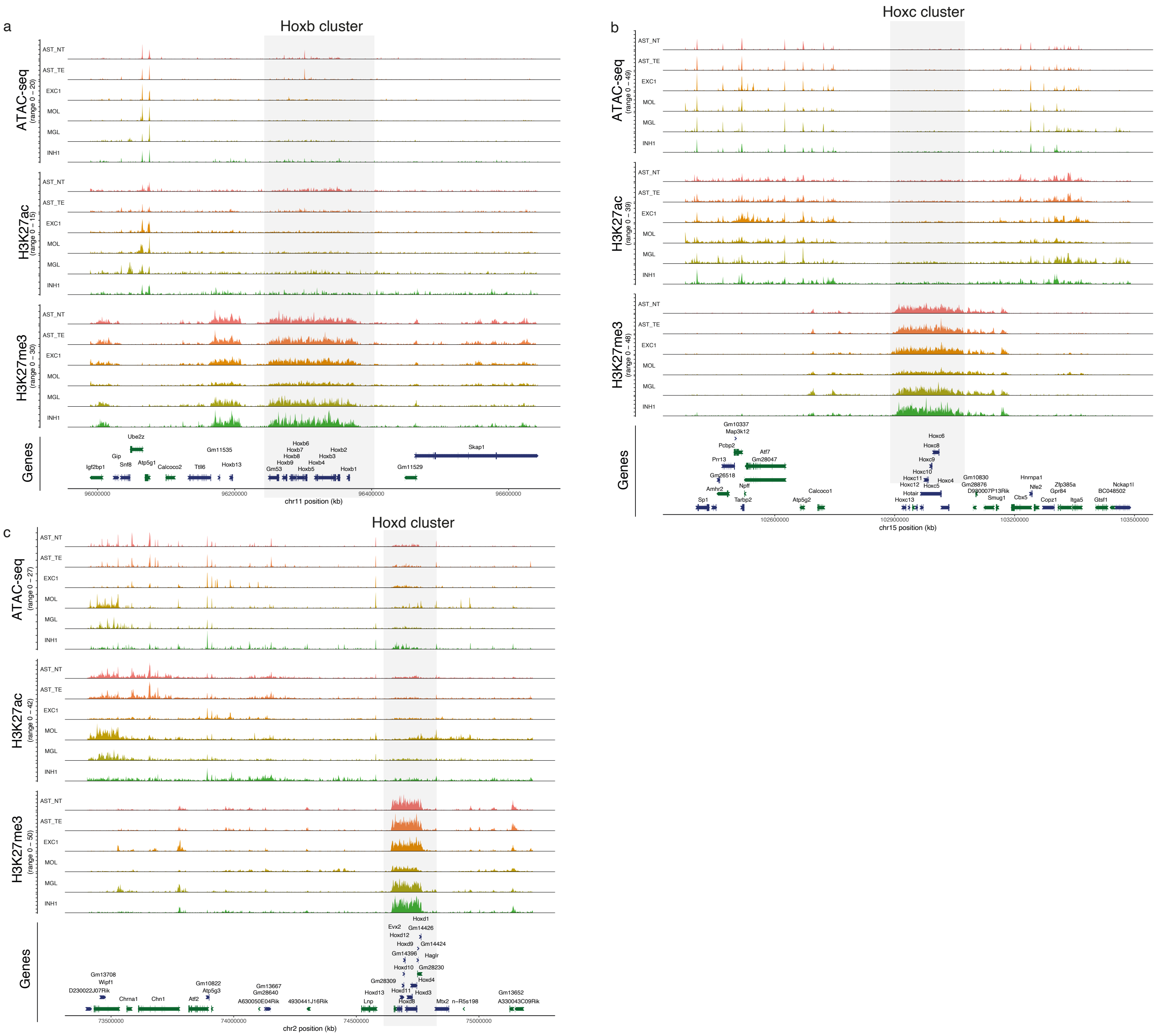

### Extended data figure 8

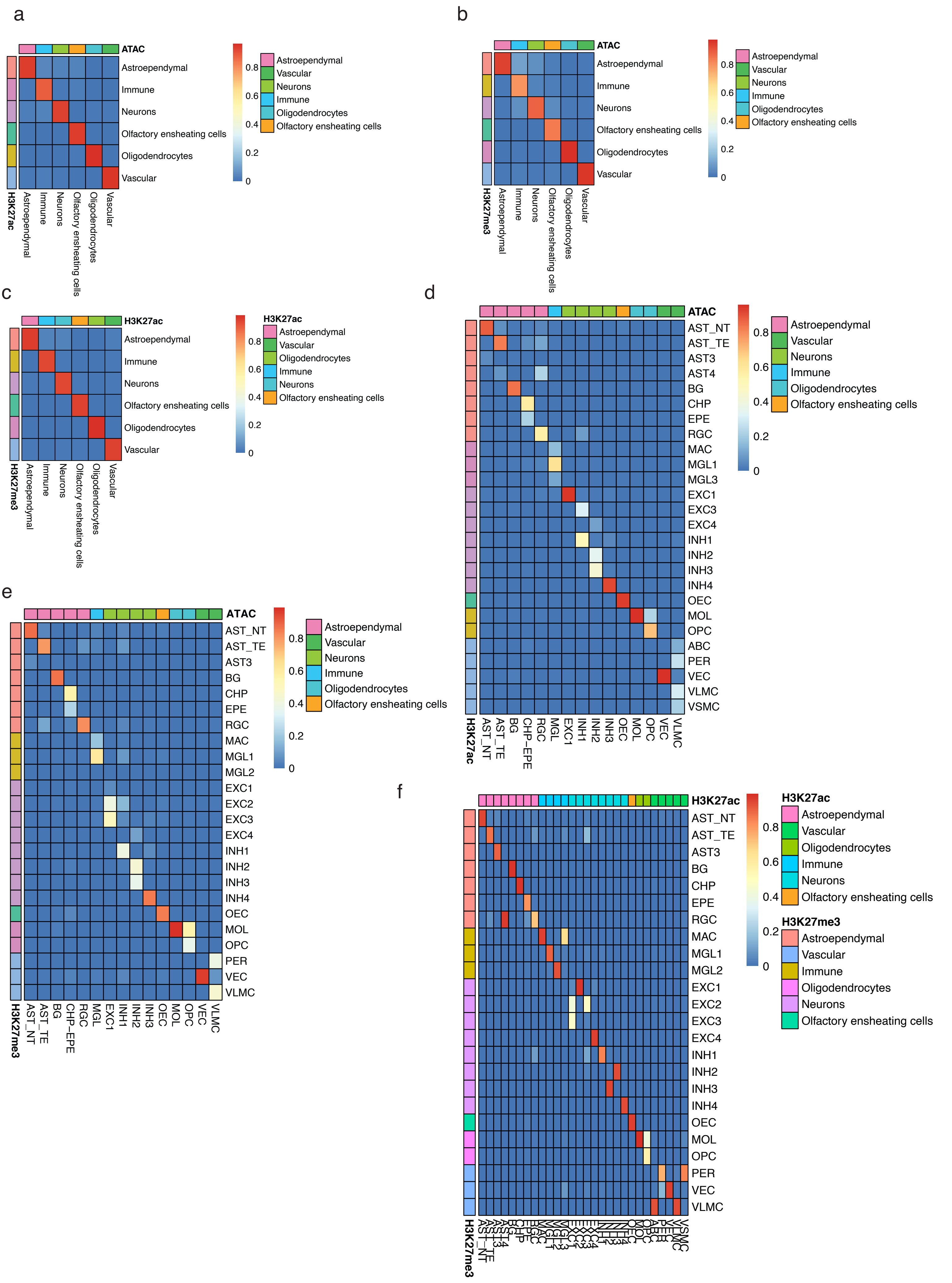

### Extended data figure 9

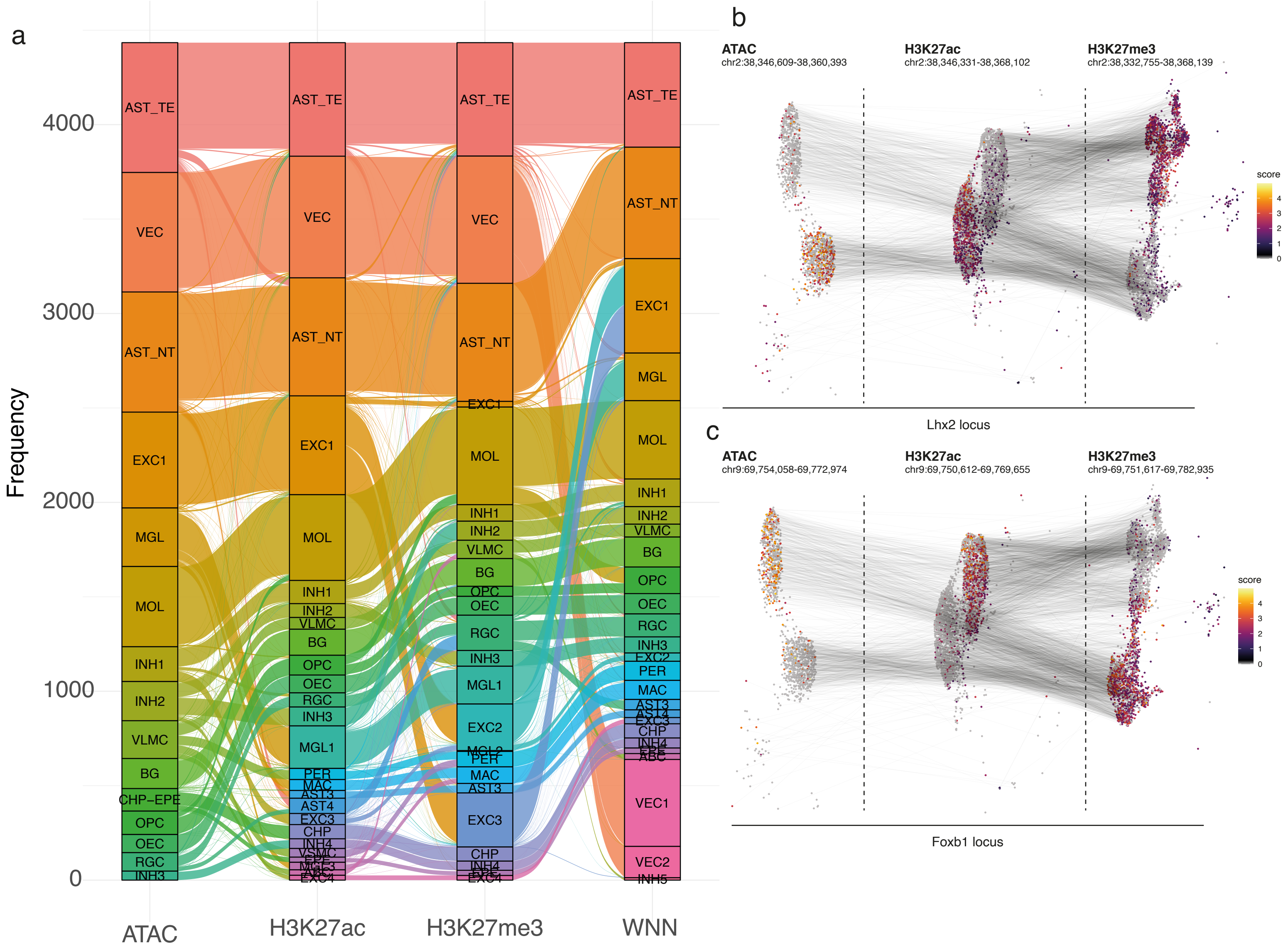

### Extended data figure 10

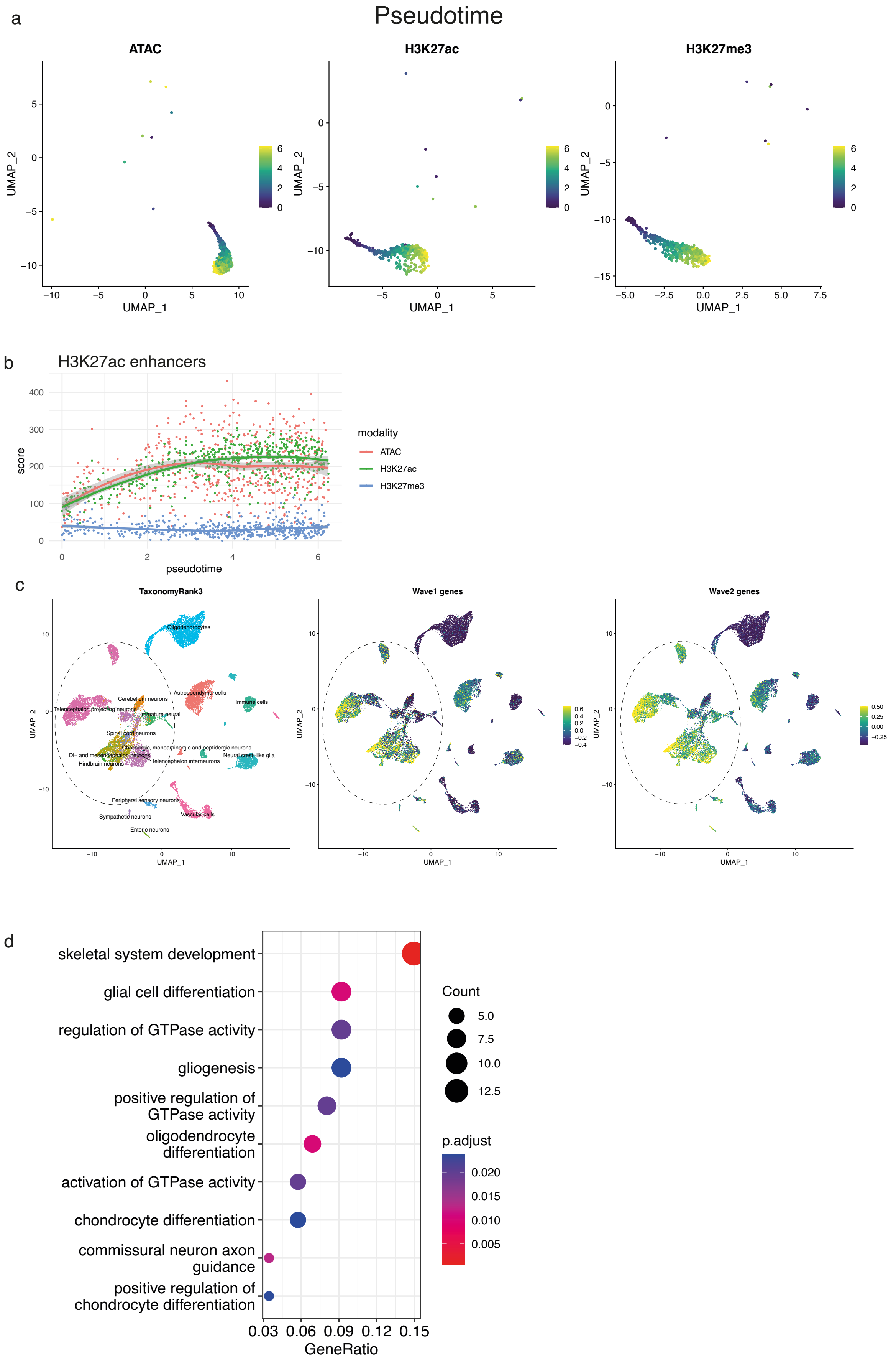

Extended data figure 11

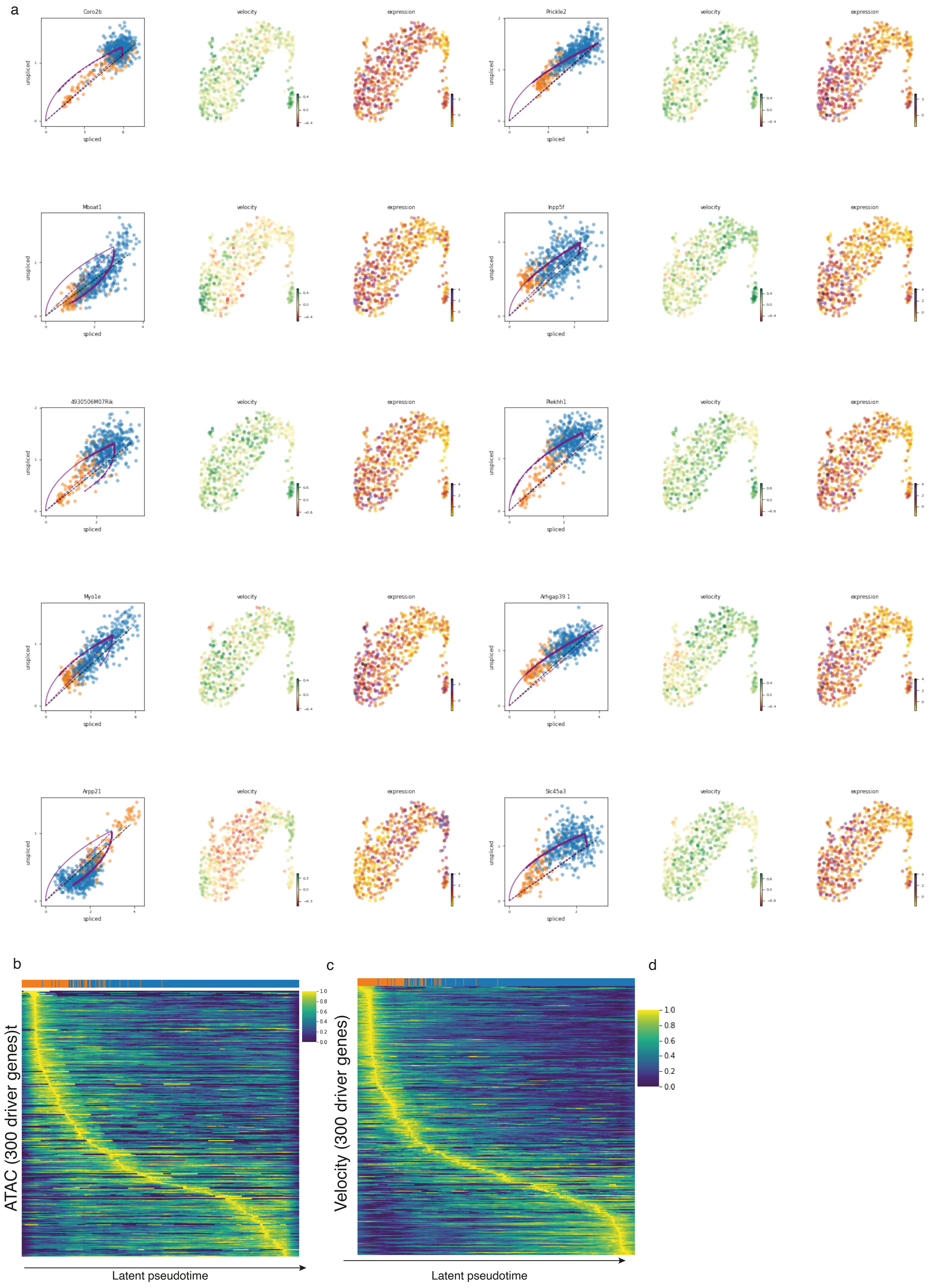
